# Identification of Potential Antibody Epitopes in MMP-15

**DOI:** 10.1101/2020.12.21.423874

**Authors:** Jessie Gan, Cheryl Eisen, Vaughn Smider

**Affiliations:** San Diego Jewish Academy, San Diego, CA 92130; Applied Biomedical Science Institute, San Diego, CA 92127; The Scripps Research Institute, Dept. of Molecular Medicine, La Jolla, CA, 92037

**Author notes:** Corresponding author: Vaughn Smider.

**Keywords:** Immunotherapy, bioinformatics, matrix metalloproteinases, epitopes, monoclonal antibody, drug development

## Abstract

Monoclonal antibody therapy is a well-established cancer treatment paradigm that often targets cancers-pecific cell surface proteins. Matrix Metalloproteinase 15 (MMP-15) is a surface protein implicated in metastasis and angiogenesis, however it is not well characterized. Here we use bioinformatics tools to identify epitopes for drug or diagnostic targeting and elucidate features of this potentially important protein in metastatic processes. We identified conserved regions in MMP-15 as well as unique, variable peptide regions compared to other Membrane Type MMPs (MT-MMP). Conservation of catalytic and hemopexin domains in MMP-15 imply functional importance, however, their similarity to other MT-MMPs discourage use as target epitopes. Our analyses also identified an MMP-15 peptide that was highly divergent from other MMPs, suggesting that it may serve as an appropriate specific epitope for a specific antibody drug. Thus, we were able to elucidate features and potential unique epitopes of MMP-15 for use in further antibody discovery and targeting.

## Introduction

Lung cancer is the number one cause of cancer-related deaths in industrialized countries. The expected 5-year survival rate in lung cancer is 15%, compared to the over 80% 5-year survival rates of breast and prostate cancers. Furthermore, 85% of diagnosed lung cancers are non-small-cell-lung cancer (NSCLC) (Tyagi) where recent advances in novel targeted therapies have not yet made a major impact on patient survival. Combinations of conventional chemotherapy, radiation, and surgery are the standard treatment for NSCLC; however, they haven’t significantly improved prognosis since 1975 (Tyagi). Immunotherapy is a novel modality that utilizes the power of the human immune system to eliminate cancer cells. Types of immunotherapies include monoclonal antibodies, antibody drug conjugates, cancer vaccines, and chimeric antigen receptor – T cells (CAR-T). These therapies function by blocking proliferation signals, recruiting lymphocytes to the tumor, developing an immune memory of tumor antigens, and targeting cytotoxins for immediate cytotoxicity (Hallam, Dotti, Ho). The linchpin of immunotherapy is the unique antigen-specificity that can distinguish between normal and cancerous cells, allowing treatment to be both potent and specific. Antigen and epitope identification is crucial to the drug development process; finding cancer-specific antigen epitopes of interest an essential challenge in the development of this therapy (Steven).

Matrix Metalloproteinases (MMPs) are zinc-dependent enzymes that proteolyze extracellular matrix (ECM). There are 24 MMPs identified in humans, rodents, and amphibians that share structural elements such as a pro-peptide of about 80 amino acids, a catalytic domain of about 170 amino acids, a linker peptide of various lengths, and a hemopexin domain of about 200 amino acids, with a few exceptions (Cathcart, Nagase). In humans, MMPs are concentrated in connective tissue and epithelial cells. Their normal function is breaking down various extracellular matrix molecules or cleaving specific proteins (Nagase, Araki).

MMP-15 is a type-1 transmembrane protein. The transmembrane MMPs (MT-MMPs) share a common structure of a signal peptide, a peptidoglycan domain, a cysteine switch, a catalytic domain, a hinge (linker 1), a hemopexin-like domain (Hpx), a stalk region (linker 2), transmembrane region, and cytoplasmic region. In addition, transmembrane MT-MMPs also have a unique insertion of 8-9 amino acids in their catalytic domain known as the MT-loop (Itoh, Araki).

Activation of TM-MMPs involve proproteins, also known as protein convertases (PCs). The sequence RXKR in the proprotein domain is the PC recognition sequence, and PCs such as furin cleave this site while the protein is still in the ER/Golgi (Itoh). The PC cleavage is dependent on the zinc binding motif in the catalytic domain and cysteine switch motif in the propeptide, which prevent proMMPs from being activated intracellularly. The zinc binding motif of HEXXHXXGXXH and the cysteine switch motif of PRCGXPD are in a Cys-Zn^2+^ coordination, preventing an essential water molecule for catalysis from binding to the zinc ion, and blocking early activation within the cell (Itoh). As the propeptide is cleaved off, this connection is severed and the zinc ion is open for catalysis as the enzyme is presented on the cell surface (Nagase, Fanjul-Fernandez).

The catalytic domain is composed of a 5-stranded β-sheet, three α-helices, and a connective loop, along with other ions for stability. There are two zinc ions (one catalytic and one structural) and up to three structural calcium ions. Among MMPs, the catalytic domains are mostly conserved with the exception of a hydrophobic S1’ pocket near the active site, which is the primary determinant of substrate specificity (Nagase, Loffek). To support the active site, the catalytic domains also contain a conserved methionine, which forms a “Met-turn” eight residues after the active domain. Aside from all MMPs, a unique feature of MT-MMP catalytic domains is an 8 amino acid loop insertion in the catalytic domain. This loop is integral for MT-MMPs’ function in the activation of proMMP-2 (Nagase, Itoh).

MMPs can regulate tumor progression processes such as cell death, proliferation, differentiation, tumor-associated angiogenesis, and malignant conversion (Loffek, Nagase, Hotary2002). As a tumor grows, it requires reorganization of its environment. Cancer fuels its need for growth through breakdown of ECM by overexpressing proteolytic enzymes like MMP-15 (Dornier). MMP-15 can degrade collagen-I, but it is not considered a major collagenolytic enzyme. However, it can also degrade fibrin and promote cellular invasion into fibrin matrices and basement membranes, the extracellular matrix that underlies all epithelia and endothelia (Hotary2002, Hotary2000, Hotary2006, Ota, Löffek). In addition, MMP-15 has been shown to proteolyze fibronectin, laminin-1, nidogen, perlecan, and NC1 as substrates, all of which are ECM components. (Itoh, D’Ortho) However, MMP-15 also contributes to cancer promotion in other ways, such as through blocking Fas and TRAIL death ligands and associating with other MMPs. (Ito, Kobayashi, Abraham, Joshi, Liu)

We are interested in therapeutic monoclonal antibodies specific to upregulated cancer antigens for tumor targeting (Pento, Aldaroish). One such possible antigen is MMP-15, which is upregulated in NSCLC and may be involved in metastasis through ECM cleavage (Magdalena, Tao). We use the cumulative information from online databanks combined with bioinformatics, the collaborative synthesis of multiple analysis tools, to further investigate the MMP-15 protein for possible specific epitopes for antibody targeting.

## Results

To assess the potentially viable epitopes of the MMP-15 protein for antibody targeting, we curated the MMP-15 sequences of several species as well as MT-MMP sequences. We employed multiple sequence alignment and analyzed various other features from the comparison such as percent identity and Shannon entropy. In addition, we utilized partial-MMP-14 crystal structures to analyze the catalytic (PDBID: 3MA2) and hemopexin (PDBID: 3C7X) domains. The overall workflow is depicted in Figure 1.

**Figure 1.**
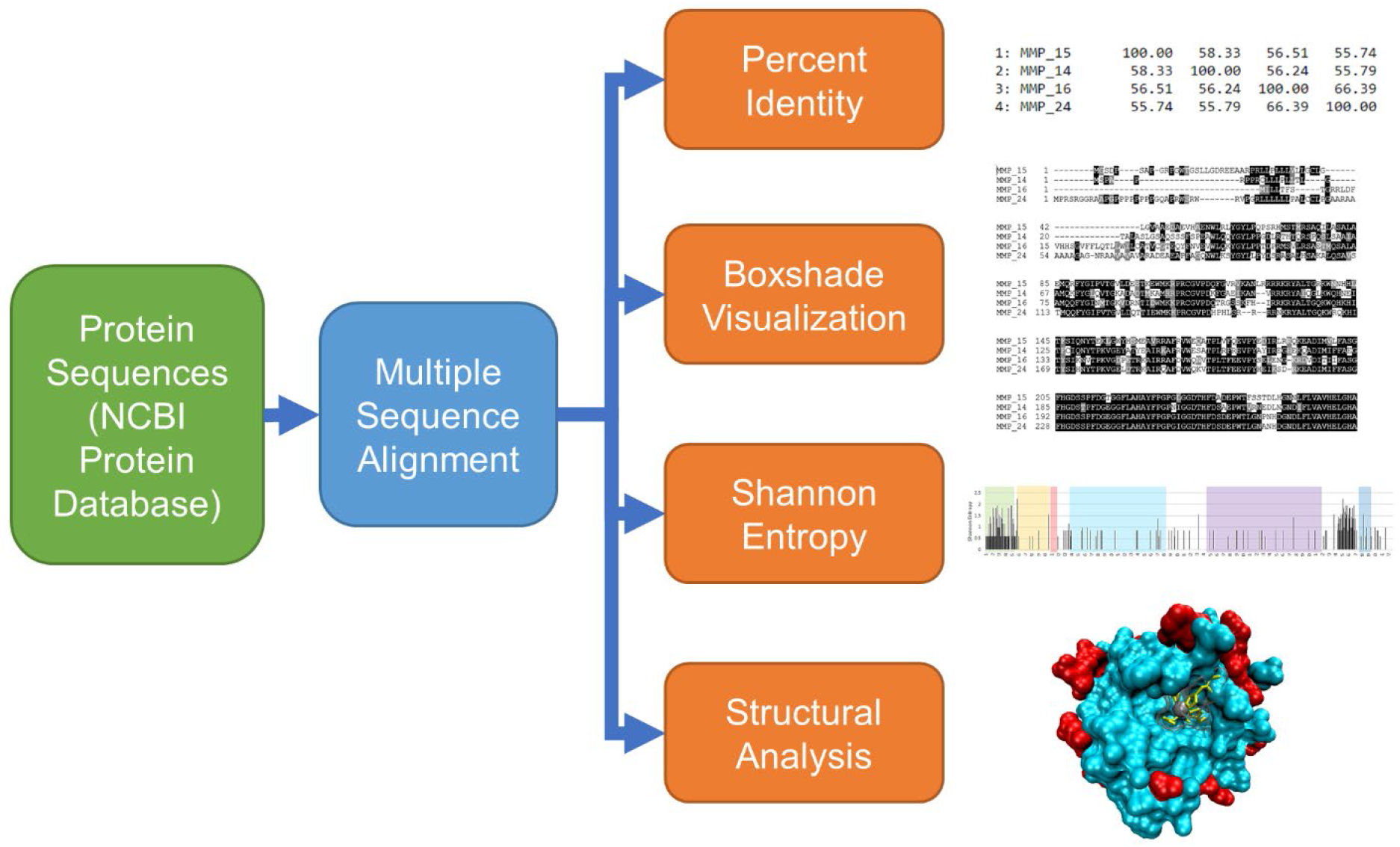
Bioinformatics workflow. Mammalian protein sequences of MMP-15 and MT-MMP sequences were obtained from the NCBI protein databank. These were aligned in Clustal Omega, and the resulting multiple sequence alignment was used in other analyses: sequence similarity metrics, BoxShade figures, Shannon entropy plots, and structural analyses.

### Mammalian MMP-15 Ortholog Comparison

We identified regions of similarity among MMP-15 sequences from various species in order to investigate regions of evolutionary conservation, which may imply their importance to functional mechanisms. Additionally, conserved epitope targeting would enable cross-reactivity of an experimental therapeutic in future animal efficacy or toxicity studies. From the multiple sequence alignment of mammalian orthologs, we found that on average 92.9% of MMP-15 amino acid residues are conserved across the aligned sequences. The enzyme active site is particularly highly conserved at an average identity of 94.7%. The sequence similarity pairwise heatmaps can be found in Supplemental Figures 1 and 2, for the entire sequence and catalytic domain respectively, and the corresponding matrixes produced from Clustal Omega are in Supplemental Tables 1 and 2.

In the mammalian MMP-15 ortholog comparison, the most sequence variability, as evaluated by Shannon entropy analysis, is seen in the signal peptide as well as a region just N-terminal to the transmembrane region (Figure 2A).

**Figure 2.**
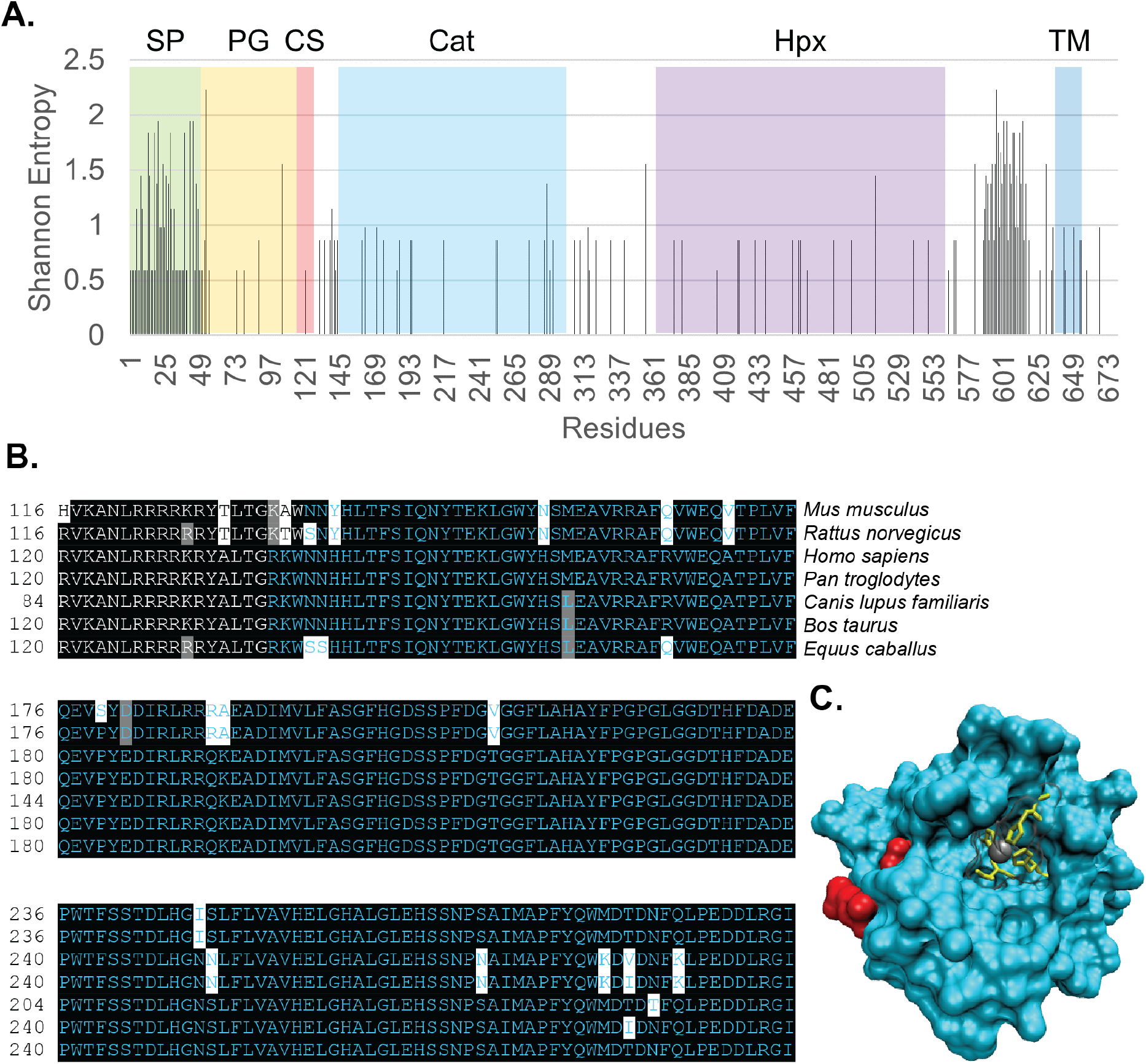
Sequence diversity of mammalian MMP-15. **A.** The mammalian MMP-15 alignment Shannon entropy shows an overall high conservation of the sequence, with the two exceptions: (i) the signal peptide (SP) and (ii) a region adjacent to the membrane. The labeled domains are the signal peptide (SP), the peptidoglycan binding domain (PG), the cysteine switch (CS), the catalytic domain (Cat), the hemopexin domains (Hpx), and the transmembrane region (TM). The Shannon entropy values for this plot can be found in Supplemental Table 3. **B.** BoxShade analysis shows high conservation of mammal MMP-15 proteins throughout the catalytic domain, colored in blue. Black highlights denote conservation, and white and grey highlights show variability. The full-length sequence BoxShade figure is provided as Supplemental Figure 3. **C.** Catalytic domain of MMP-14 (PDB: 3MA2) in surface representation shows relative conservation. The yellow active site is composed of histidines that hold a zinc ion (grey) that is a feature of MMP’s in their proteolytic mechanism. The variable residues for a Shannon entropy greater than 0.5 for the MMP-15 mammalian alignment are highlighted in red.

As shown in Figure 2B and 2C, the active site of the catalytic domain is highly conserved. Although there are a few variable residues in the catalytic domain, they are on the opposite side of the conserved active site. The two visible variable residues in Figure 2C are N114 and T78 (SE> 0.5); all variable residues in the catalytic domain had low Shannon entropy scores near 1, indicating considerable conservation. In addition, structural analysis of the hemopexins additionally showed that this is a highly conserved domain (Supplemental Figure 4).

### Membrane-Type MMP Paralog Comparison

In order to identify regions within MMP-15 that are unique compared to other MT-MMP’s, we compared four human MT-MMP (MMP-14, 15, 16, and 24) paralog sequences in a multiple sequence alignment. We found that on average 56.9% of residues are conserved among the MMPs compared. The enzyme active site is more conserved at an average of 68.5%. The sequence similarity heatmap across the entire sequence is provided in Supplemental Figure 5, with the corresponding matrix from Clustal Omega in Supplemental Table 4.

As there is currently no structural information on MMP-15 itself, we used the crystal structure of the related MMP-14 catalytic domain as a surrogate to visualize structural diversity. The catalytic domain has clear areas of structural conservation, as well as more variable regions than the mammalian alignment (Figure 3). However, the majority of the variable residues are on the side opposite the active site, leaving the active site itself, and residues surrounding it, mostly conserved. Thus, the face of the enzyme containing the active stie is highly conserved whereas the opposite face appears much more mutable. In addition, compared to the hemopexin domains in the mammalian MMP-15 alignment, the hemopexin domains of the multiple MT-MMP alignment are also much more variable (Supplemental Figure 7).

**Figure 3.**
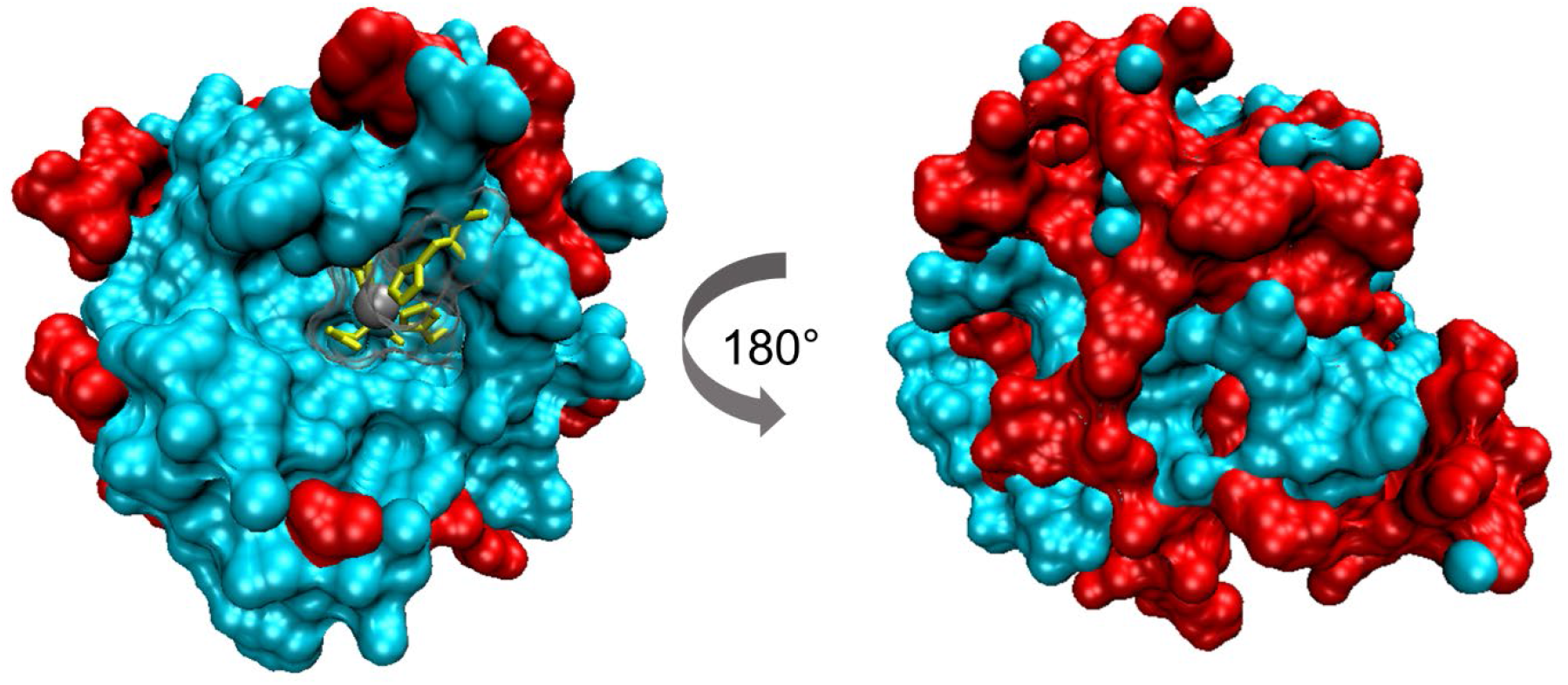
Structural diversity of MMP-15 catalytic domain compared to other MT-MMPs. The MT-MMP alignment reveals conserved and variable regions of the catalytic. The catalytic domain of crystal MMP-14 is in surface representation. The yellow active site is composed of histidines that hold a zinc ion (grey) that is a feature of MMP’s in their proteolytic mechanism. The variable residues of the MT-MMP alignment based on Shannon entropy values ranging from 0.8 to 1.5 are highlighted in red.

According to the Shannon entropy results (Figure 4), the MT-MMP paralog alignment shows more variability than the mammalian alignment of MMP-15 orthologs. In two select areas within the catalytic domain of paralogs, there is nearly a complete conservation of residues, whereas in other domains there is no such conservation. In addition to revealing the most conserved regions, Shannon entropy analysis also identifies the most variable regions, which may be of interest as novel antibody targeting epitopes.

**Figure 4.**
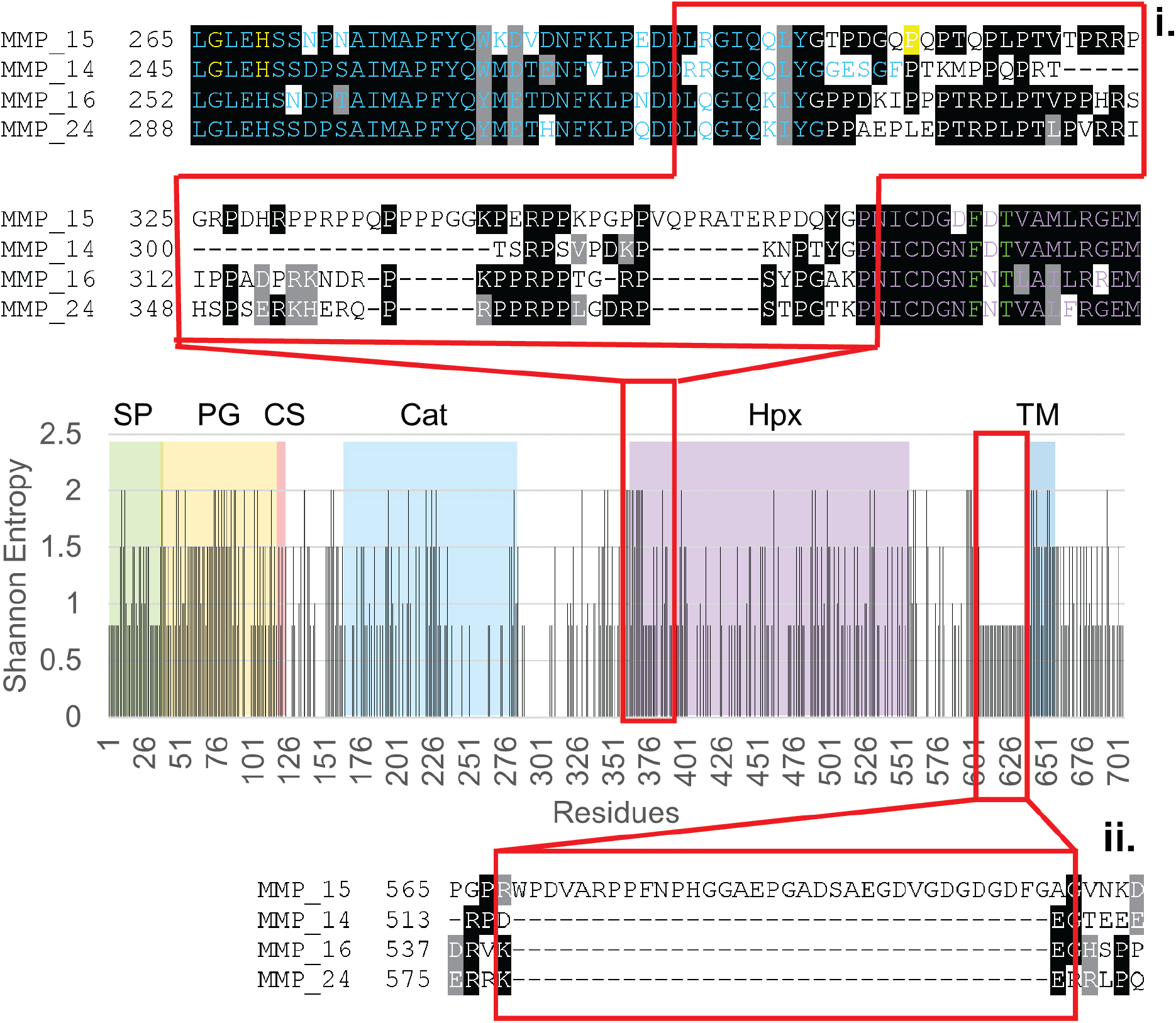
Variable regions of the MT-MMP paralogs. The MT-MMP paralog alignment reveals variable regions for possible antibody targeting as well as conserved catalytic domains. The Shannon entropy values for this plot can be found in Supplemental Table 5, and they are enumerated by the full MT-MMP alignment. Sequence **i.** is residues 310 to 343 (relative to the MMP-15 sequence) and is highly variable. Sequence **ii.** is residues 569 to 603, which is an insertion that is completely unique to MMP-15. The full sequence BoxShade figure is provided in Supplemental Figure 6.

Specifically, two sequence regions, labeled as sequences i. and ii. in Figure 4, are shown in BoxShade alignments and have significant variability between paralogs. The two regions are amino acid residues 310 to 343, and 569 to 603. The first region, residues 310-343, is in a proline-rich linker region between the catalytic and hemopexin domain extending into the N-terminal portion of the hemopexin domain. Residues 569-603 are a stretch of 34 amino acids that are completely unique to MMP-15 compared to other MT-MMP family members. This sequence appears to result from an insertion event in MMP-15 which did not occur in other MT-MMPs.

## Discussion

In the development of cancer therapeutics, drug specificity has become integral to improving upon current treatments, as non-specific therapies may have significant toxicity. Immunotherapy utilizes the specificity of the immune system to target cancer cells, and the potential for identifying the most optimal target antigens and epitopes is increasingly important. In this work we investigate two comparisons of the potential antibody target MMP-15: i) MMP-15 orthologous sequences (across species) and ii) human MT-MMP paralogous sequences, in order to identify conserved, and thereby likely important residues and to identify novel, unique epitopes of MMP-15, respectively. Sequence regions that are both essential to function and unique to the protein of interest have potential as epitopes for effective drug targeting, to impair target function and reduce toxicity, respectively. Conserved residues across MMP-15 orthologs, but divergent across MT-MMP paralogs, would not only optimize drug development by targeting residues conserved in function, but also may enable non-human efficacy and toxicity studies while minimizing off-target effects by limiting binding to paralog epitopes. Here we identify such regions and suggest their viability as target epitopes.

In the mammalian MMP-15 ortholog analysis, the linker region before the transmembrane region is unannotated as a protein domain and is relatively variable. In addition, the high conservation of the MMP-15 catalytic domain suggests that the MMP-15 substrate specificity between mammals is likely preserved. The docking of the substrate is dictated by the S1 pocket which varies in depth among MMPs and is the binding site for substrates (Cathcart, Nagase). In addition, our results support hemopexins as an important domain through their high conservation (Itoh, Butler). Previous work with MMP-14 demonstrated that the hemopexin domains are used for dimerization of MMP’s, and hemopexins appear to be present in all MMPs (Nagase).

From the MT-MMP paralog analysis, we have identified key features across MT-MMP’s as well as possible targeting regions with significant variability. There is an even distribution of variation across the entire sequence, however the placement of variation and the domains they are in may have significance. Not surprisingly, the catalytic domain has higher conservation, due to its importance in performing its function. The more variable regions of the catalytic domain may contain residues that control substrate specificity through altering the surface of the catalytic domain, such as the hydrophobic S1’ pocket. Although these MT-MMP’s are of the same family with their common transmembrane region, they may have diverse substrates (Nagase). The hemopexin domains were more diverse within the MT-MMP paralog comparison than the mammalian ortholog comparison, likely due to the fact that different types of MT-MMPs may have evolved divergent functions, and the hemopexin domain itself may have evolved unique dimerization components.

The aim of this work is to identify possible epitope sequences. Variable regions between MT-MMPs is relevant to identifying more specific epitopes. In studies of MMP-14, complications of previous MMP small molecule inhibitors included unpredicted side effects due to the off-target inhibition of other MMPs (Overall, Giavazzi). If an antibody was directed to target highly conserved regions of paralogs, then it could bind to any MT-MMP and have off-target consequences. Therefore, regions that are unique to MMP-15 (versus other MMPs) are the potentially optimal epitopes. We identified residues 310-343 as highly variable, with high divergence from other sequences that might yield higher specificity of the antibody to MMP-15 only, lowering possible off-target toxicities. However, residues 569-603 appear to result from an insertional event and are completely distinctive to MMP-15, implying even higher potential specificity. This peptide appears to have a long extension which may allow MMP-15 a wider range of mobility on the cell surface than other MMPs. Alternatively, it may have an as yet undiscovered structure and function. Of considerable importance, these two regions may provide for optimal epitopes for antibody targeting. In this regard, high affinity, specific antibodies to these regions could be useful as therapeutics, diagnostics, or valuable research reagents.

Our results reveal that open-source bioinformatics tools could be used to identify conserved and diverse regions in the MMP-15 protein compared to other MMP-15 orthologs and other MT-MMP paralogs. These regions can be considered as potentially specific antigen epitopes of MMP-15 in consideration for antibody, drug, or diagnostic targeting. The next step is to use the sequences identified to discover and engineer high affinity monoclonal antibodies experimentally, which could be used in NSCLC or other diseases as therapeutics or diagnostics.

## Experimental Procedures

We analyzed MMP-15 orthologous amino acid sequences from *Homo sapiens* (NP_002419.1)*, Mus musculus* (NP_032635.1), *Rattus norvegicus* (NP_001099638.1), *Bos taurus* (NP_001178363.1)*, Canis lupus familiaris* (XP_022270500.1)*, Pan troglodytes* (XP_009429160.2)*, Equus caballus* (XP_023492845.1). In a separate alignment, we analyzed four paralogous *Homo Sapiens* type I MT-MMPs: MMP-15, MMP-14 (NP_004986.1), MMP-16 (NP_005932.2), and MMP-24 (NP_006681.1), in the transmembrane (TM) paralog alignment. The amino acid sequences for both comparisons were procured from NCBI’s protein data base. Both comparisons were aligned using Clustal Omega (Clustal), an online multiple sequence alignment software, which can be found at https://www.ebi.ac.uk/Tools/msa/clustalo/. The program inputs were the FASTA sequences from NCBI (NCBI), and the output was specified as a Pearson/FASTA file. The results from Clustal Omega were also displayed with BoxShade using the BoxShade server at https://embnet.vital-it.ch/software/BOX_form.html (BoxShade).

From the alignments, a variability plot was also performed using the Protein Variability Server at http://imed.med.ucm.es/PVS/, which calculates variability per residue (PVS). The variability plots were constructed for both the mammalian ortholog comparison and the MT-MMP paralog comparison through the Shannon entropy method, however for the orthologs the reference sequence was the consensus sequence while for the MT-MMP paralog comparison MMP-15 was the reference sequence. The Shannon entropy analysis is considered one of the most sensitive methods for estimating the diversity of a system (Shannon, Litwin). Additional plots were made for the variability of the catalytic domains and the hemopexin domains for both analyses with the same selections.

After analyzing the sequences, we performed a structural analysis in the visualization program, Visual Molecular Dynamics (VMD) (Humphrey). Using the protein annotations from NCBI, domains were projected onto the 3-D structures according to residue. In addition, variable residues were investigated and visualized as described in each figure. The structures for both the mammalian species ortholog comparison and the TM-MMP paralog comparison are portions of various crystal structures of MMP-14, pdbs 3MA2 for the catalytic domain and 3C7X for the hemopexin domains. Both crystal structures were from RCSB PDB (https://www.rcsb.org/), an online database for protein crystal structures (Berman, Grossman, Tochowicz). MMP-14 was used as a surrogate for MMP-15 (57.9% identity) because there is currently no available crystal structure of MMP-15.

