## Supplementary Material for "Identification of Potential Antibody Epitopes in MMP-15"

### Supplemental Materials

#### a. Figures

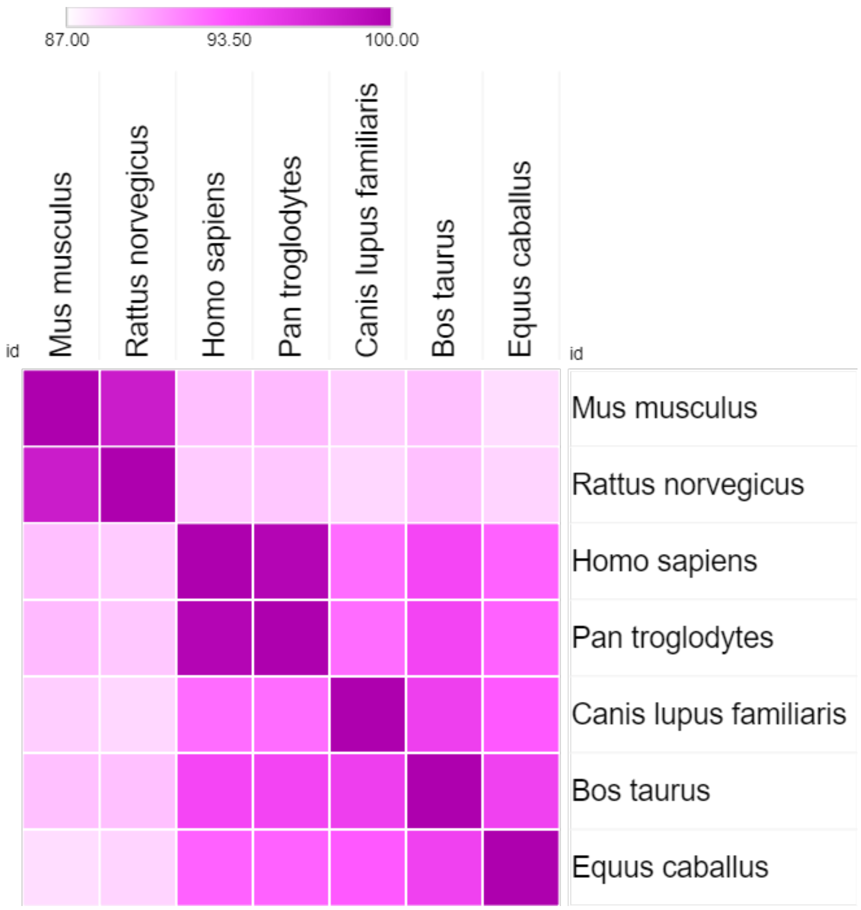


Supplemental Figure 1. Mammalian MMP-15 ortholog percent identity heatmap over the entire sequence. The percent identity matrix was produced with Clustal omega, (see Supplemental Table 1). The heatmap was generated in the Morpheus online heatmap generator at https://software.broadinstitute.org/morpheus/.

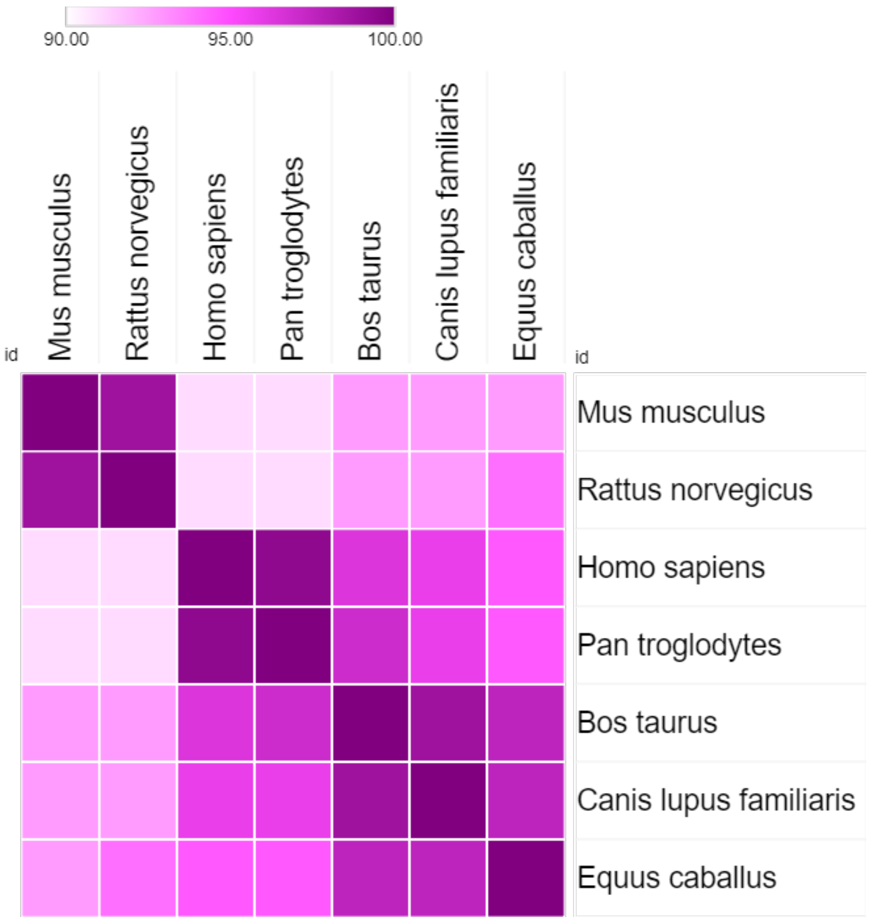


Supplemental Figure 2. Mammalian MMP-15 catalytic domain percent identity heatmap. The percent identity similarity matrix was produced with Clustal Omega (see Supplemental Table 2). The heatmap was generated in the Morpheus online heatmap generator at https://software.broadinstitute.org/morpheus/.

```
Mus_muscul     1 MGSDRSALGRPGCTGSCLSS-----RASLLPLLLVLLDCLGHGTASKDAEVYAAENWLRL
Rattus_nor     1 MGSDRSALGRPGCAGSCLSS-----RASLLLLLLVLLDCLGGGTASEDAEVYAAENWLRL
Homo_sapie     1 MGSDPSAPGRPGWTGSLLGDREEAARPRLLPLLLVLLGCLGLGVAAEDAEV-HAENWLRL
Pan_troglo     1 MGSDRSAPGRPGWTGSVLGDREEAARPRLLPLLLVLLGCLGLGVAAEDAEV-HAENWLRL
Canis_lupu     1 -----------------------------------MVTAVGGG-SGPNRAV-VAVNWLRL
Bos_taurus     1 MGSDRSAPGRPGWAGSLLGGREAATRPQLLPLLLVLLSCLGRGAAAEDAEV-NAENWLRL
Equus_caba     1 MGSDRSALARPGRAGSLLGGREAAARPRMLPLLLVLLGCLGRGAAAEDAEV-KAENWLRL

Mus_muscul    56 YGYLPQPSRHMSTMRSAQILASALAEMQSFYGIPVTGVLDEETKTWMKRPRCGVPDQFGV
Rattus_nor    56 YGYLPQPSRHMSTMRSAQILASALAEMQSFYGIPVTGVLDEETKTWMKRPRCGVPDQFGV
Homo_sapie    60 YGYLPQPSRHMSTMRSAQILASALAEMQRFYGIPVTGVLDEETKEWMKRPRCGVPDQFGV
Pan_troglo    60 YGYLPQPSRHMSTMRSAQILASALAEMQRFYGIPVTGVLDEETKEWMKRPRCGVPDQFGV
Canis_lupu    24 YGYLPQPSRHMSTMRSAQILASALAEMQRFYGIPVTGVLDEETKAWMKRPRCGVPDQFGV
Bos_taurus    60 YGYLPQPSRHMSTMRSAQILASALAEMQRFYGIPVTGVLDEETKAWMKRPRCGVPDQFGV
Equus_caba    60 YGYLPQPSRHMSTLRSAQTLASALAEMQRFYGIPVTGVLDEETKAWMKRPRCGVPDQFGV

Mus_muscul   116 HVKANLRRRRKRYTLTGKAWNNYHLTFSIQNYTEKLGWYNSMEAVRRAFQVWEQVTPLVF
Rattus_nor   116 RVKANLRRRRRRYTLTGKTWSNYHLTFSIQNYTEKLGWYNSMEAVRRAFQVWEQVTPLVF
Homo_sapie   120 RVKANLRRRRKRYALTGRKWNNHHLTFSIQNYTEKLGWYHSMEAVRRAFRVWEQATPLVF
Pan_troglo   120 RVKANLRRRRKRYALTGRKWNNHHLTFSIQNYTEKLGWYHSMEAVRRAFRVWEQATPLVF
Canis_lupu    84 RVKANLRRRRKRYALTGRKWNNHHLTFSIQNYTEKLGWYHSLEAVRRAFRVWEQATPLVF
Bos_taurus   120 RVKANLRRRRKRYALTGRKWNNHHLTFSIQNYTEKLGWYHSLEAVRRAFRVWEQATPLVF
Equus_caba   120 RVKANLRRRRRRYALTGRKWSSHHLTFSIQNYTEKLGWYHSLEAVRRAFQVWEQATPLVF

Mus_muscul   176 QEVSYDDIRLRRRAEADIMVLFASGFHGDSSPFDGVGGFLAHAYFPGPGLGGDTHFDADE
Rattus_nor   176 QEVPYDDIRLRRRAEADIMVLFASGFHGDSSPFDGVGGFLAHAYFPGPGLGGDTHFDADE
Homo_sapie   180 QEVPYEDIRLRRQKEADIMVLFASGFHGDSSPFDGTGGFLAHAYFPGPGLGGDTHFDADE
Pan_troglo   180 QEVPYEDIRLRRQKEADIMVLFASGFHGDSSPFDGTGGFLAHAYFPGPGLGGDTHFDADE
Canis_lupu   144 QEVPYEDIRLRRQKEADIMVLFASGFHGDSSPFDGTGGFLAHAYFPGPGLGGDTHFDADE
Bos_taurus   180 QEVPYEDIRLRRQKEADIMVLFASGFHGDSSPFDGTGGFLAHAYFPGPGLGGDTHFDADE
Equus_caba   180 QEVPYEDIRLRRQKEADIMVLFASGFHGDSSPFDGTGGFLAHAYFPGPGLGGDTHFDADE

Mus_muscul   236 PWTFSSTDLHGISLFLVAVHELGHALGLEHSSNPSAIMAPFYQWMDTDNFQLPEDDLRGI
Rattus_nor   236 PWTFSSTDLHGISLFLVAVHELGHALGLEHSSNPSAIMAPFYQWMDTDNFQLPEDDLRGI
Homo_sapie   240 PWTFSSTDLHGNNLFLVAVHELGHALGLEHSSNPNAIMAPFYQWKDVDNFKLPEDDLRGI
Pan_troglo   240 PWTFSSTDLHGNNLFLVAVHELGHALGLEHSSNPNAIMAPFYQWKDIDNFKLPEDDLRGI
Canis_lupu   204 PWTFSSTDLHGNSLFLVAVHELGHALGLEHSSNPSAIMAPFYQWMDTDTFQLPEDDLRGI
Bos_taurus   240 PWTFSSTDLHGNSLFLVAVHELGHALGLEHSSNPSAIMAPFYQWMDIDNFQLPEDDLRGI
Equus_caba   240 PWTFSSTDLHGNSLFLVAVHELGHALGLEHSSNPSAIMAPFYQWMDTDNFQLPEDDLRGI

Mus_muscul   296 QQLYGSPDGKPQPTRPLPTVRPRRPGRPDHQPPRPPQPPHPGGKPERPPKPGPPPQPRAT
Rattus_nor   296 QQLYGSPDGKPQPTRPLPTVRPRRPGRPDHQPPRPPQPPHPGGKPERPPKPGPPPQPRAT
Homo_sapie   300 QQLYGTPDGQPQPTQPLPTVTPRRPGRPDHRPPRPPQPPPPGGKPERPPKPGPPVQPRAT
Pan_troglo   300 QQLYGTPDGQPQPTQPLPTVTPRRPGRPDHRPPRPPQPPPPGGKPERPPKPGPPVQPRAT
Canis_lupu   264 QQLYGTPDGQPQPTRPLPTVTPRRPGRPDHRPPRPPQPPPPGGKPERPPKPGPPAQPRAT
Bos_taurus   300 QQLYGTPDGQPQPTQPLPTVTPRRPGRPDHRPPRPPQPPPPGGKPERPPKPGPPPQPRAT
Equus_caba   300 QQLYGTPDGQPQPTRLLPTVTPRRPGRPDHRPPRPPQPPPPGGKPERPPKPGPPAQPRAT
```

```
Mus_muscul   356 ERPDQYGPNICDGNFDTVAVLRGEMFVFKGRWFWRVRHNRVLDNYPMPIGHFWRGLPGNI
Rattus_nor   356 ERPDQYGPNICDGNFDTVAVLRGEMFVFKGRWFWRVRHNRVLDNYPMPIGHFWRGLPGNI
Homo_sapie   360 ERPDQYGPNICDGDFDTVAMLRGEMFVFKGRWFWRVRHNRVLDNYPMPIGHFWRGLPGDI
Pan_troglo   360 ERPDQYGPNICDGDFDTVAMLRGEMFVFKGRWFWRVRHNRVLDNYPMPIGHFWRGLPGDI
Canis_lupu   324 ERPDQYGPNICDGDFDTVAMLRGEMFVFKGRWFWRVRHNRVLDNYPMPIGHFWRGLPSDI
Bos_taurus   360 ERPDQYGPNICDGDFDTVAMLRGEMFVFKGRWFWRVRHNRVLDNYPMPIGHFWRGLPSDI
Equus_caba   360 ERPDQYGPNICDGDFDTVAMLRGEMFVFKGRWFWRVRHNRVLDSYPMPIGHFWRGLPGDI

Mus_muscul   416 SAAYERQDGHFVFFKGNRYWLFREANLEPGYPQPLSSYGTDIPYDRIDTAIWWEPTGHTF
Rattus_nor   416 SAAYERQDGHFVFFKGNRYWLFREANLEPGYPQPLSSYGTDIPYDRIDTAIWWEPTGHTF
Homo_sapie   420 SAAYERQDGRFVFFKGDRYWLFREANLEPGYPQPLTSYGLGIPYDRIDTAIWWEPTGHTF
Pan_troglo   420 SAAYERQDGRFVFFKGDRYWLFREANLEPGYPQPLTSYGLGIPYDRIDTAIWWEPTGHTF
Canis_lupu   384 SAAYERQDGRFVFFKGDRYWLFREANLEPGYPQPLTSYGLGIPYDRIDTAIWWEPTGHTF
Bos_taurus   420 SAAYERQDGRFVFFKGDRYWLFREANLEPGYPQPLTSYGLGIPYDRIDTAIWWEPTGHTF
Equus_caba   420 SAAYERQDGRFVFFKGDRYWLFREANLEPGYPQPLTSYGLGIPYDHIDTAIWWEPTGHTF

Mus_muscul   476 FFQADRYWRFNEETQHGDPGYPKPISVWQGIPTSPKGAFLSNDAAYTYFYKGTKYWKFNN
Rattus_nor   476 FFQADRYWRFNEETQHGDPGYPKPISVWQGIPNSPKGAFLSNDAAYTYFYKGTKYWKFNN
Homo_sapie   481 FFQEDRYWRFNEETQRGDPGYPKPISVWQGIPASPKGAFLSNDAAYTYFYKGTKYWKFDN
Pan_troglo   481 FFQEDRYWRFNEETQRGDPGYPKPISVWQGIPASPKGAFLSNDAAYTYFYKGTKYWKFDN
Canis_lup    445 FFQEDRYWRFNEETQRGDPGYPKPISVWQGIPTSPKGAFLSNDAAYTYFYKGTKYWKFDN
Bos_taurus   481 FFQEDRYWRFNEETQRGDPGYPKPISVWQGIPASPKGAFLSNDAAYTYFYKGTKYWKFDN
Equus_caba   481 FFQEDRYWRFNEETQRGDPGYPKPISVWQGIPTSPKGAFLSNDAAYTYFYKGTKYWKFDN

Mus_muscul   537 ERLRMEPGHPKSILRDFMGCQEHVEPRSRWPDVARPPFNPNGGAEPEADGDSKEENA---
Rattus_nor   537 ERLRMEPGHPKSILRDFMGCQEHVEPRSRWPDVARPPFNPNGGAEPETEDDSKEEDV---
Homo_sapie   541 ERLRMEPGYPKSILRDFMGCQEHVEPGPRWPDVARPPFNPHGGAEPGADSA--------E
Pan_troglo   541 ERLRMEPGYPKSILRDFMGCQEHVEPGPRWPDVARPPFNPHGGAEPGADSA--------E
Canis_lup    505 ERLRMEPGYPKSILRDFMGCQEQVEPGPRWPDVARPPFNPDGGAEPGAGGDGEEGEEGHE
Bos_taurus   541 ERLRMEPGYPKSILRDFMGCQEHVEPGPRWPDVARPPFNPDGGAEPGMGGDSEEGEEGGE
Equus_caba   541 ERLRMEPGYPKSILRDFMGCQEHVEPGPRWPDVARPPFNPDGGAEPGAGGDSEEGSEGPA

Mus_muscul   581 -------------GDKDEGSRVVVQMEEVVRTVNVVMVLVP-LLLLLCILGLAFALVQMQ
Rattus_nor   581 -------------GDKDEGSRVVVQMEEVVRTVNVVMVLVP-LLLLLCILGLAFALVQMQ
Homo_sapie   593 GDVGDGDGDFGAGVNKDGGSRVVVQMEEVARTVNVVMVLVP-LLLLLCVLGLTYALVQMQ
Pan_troglo   593 GDVGDGDGDFGAGVNKDGGSRVVVQMEEVARTVNVVMVLVP-LLLLLCVLGLTYALVQMQ
Canis_lup    565 PGTGGRDQDSGEDTDEDGGSRVVVQLEEVTRTVNMVMVLVPPLLLLLCILGLTYALVQMQ
Bos_taurus   601 AGPGGGGE---AGPDEDGGSRVVVQMEEVTRTVNMVMVLVPPVLLLLCILGLTYALVQMQ
Equus_caba   601 AGPGGGEGDFGVGTDGDGGSRVVVQMEEVTRTVSMVMVLVPPLLLLLCVLGLTYALVHMQ

Mus_muscul   627 RKGAPRMLLYCKRSLQEWV
Rattus_nor   627 RKGAPRMLLYCKRSLQEWV
Homo_sapie   652 RKGAPRVLLYCKRSLQEWV
Pan_troglo   652 RKGAPRVLLYCKRSLQEWV
Canis_lup    625 RKGAPRMLLYCKRSLQEWV
Bos_taurus   658 RKGAPRMLLYCKRSLQEWV
Equus_caba   661 RKGAPRVLLYCKRSLQEWV
```

Key:
Light Green – Signal Peptide
Orange – Putative peptidoglycan binding domain
Red – Cysteine Switch (MMP14 is not annotated, but highly conserved)
Dark Green – Proprotein convertase recognition sequence (Nagase)
Teal – Catalytic Domain (Peptidase M10)
Yellow – Active Site (MMP16 & 24 not annotated, but highly conserved)
Hemopexin 1 Hemopexin 2 Hemopexin 3 Hemopexin 4 – Hemopexin Domains
Bright Green – Metal Binding

Supplemental Figure 3. Boxshade alignment of mammalian MMP-15 orthologs, labeled by color according to NCBI annotations.

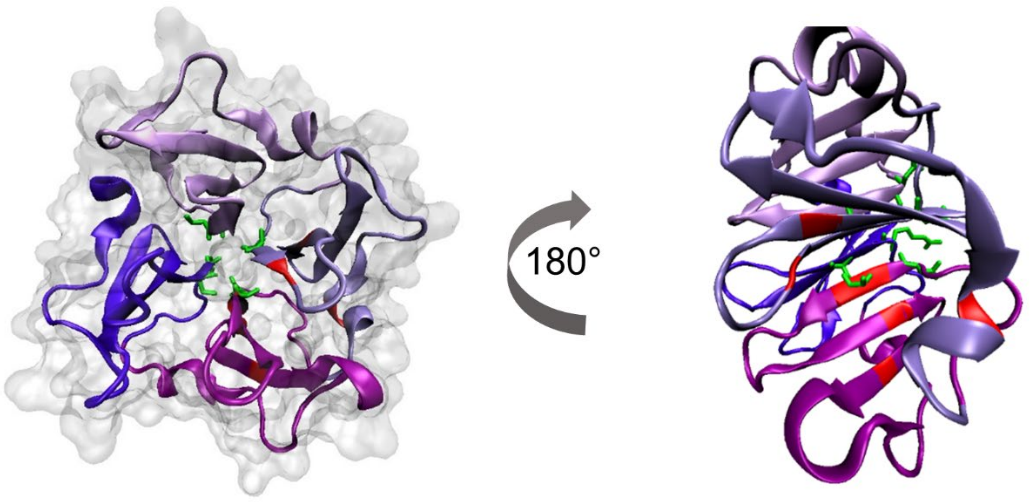


Supplemental Figure 4. Hemopexin domains of MMP-14 (as a surrogate for MMP-15) in surface and new cartoon representation show very little variability in the mammalian alignment. The metal binding sites are represented in green and variable regions are represented in red. In the right figure, the hemopexin domain is rotated to show areas of variability within hemopexin 2 and 3.

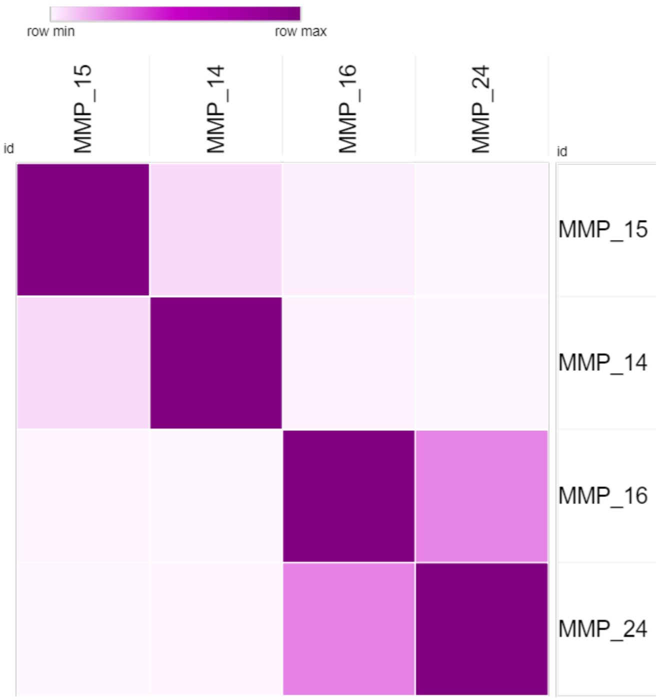


Supplemental Figure 5. MT-MMP paralog percent identity heatmap over the entire sequence. The percent identity matrix was produced with Clustal Omega (see Supplemental Table 3). The heatmap was generated in the Morpheus online heatmap generator at https://software.broadinstitute.org/morpheus/.

```
MMP_15     1 --------MGSDP----SAP-GRPGWTGSLLGDREEAARPRLLPLLLVLLGCLG------
MMP_14     1 --------MSPA----P--------------------RPPRCLLLPLLTL----G-----
MMP_16     1 -----------------------------------------MILLTFS-----TGRRLDF
MMP_24     1 MPRSRGGRAAPGPPPPPPPPGQAPRWSRW-------RVPGRLLLLLLPALCCLPGAARAA

MMP_15    42 -----------------LGVAAEDAEVHAENWLRLYGYLPQPSRHMSTMRSAQILASALA
MMP_14    20 -------------TALASLGSAQSSSFSPEAWLQQYGYLPPGDLRTHTQRSPQSLSAAIA
MMP_16    15 VHHSGVFFLQTLLWILCATVCGTEQYFNVEVWLQKYGYLPPTDPRMSVLRSAETMQSALA
MMP_24    54 AAAAGAG-NRAAVAVAVARADEAEAPFAGQNWLKSYGYLLPYDSRASALHSAKALQSAVS

MMP_15    85 EMQRFYGIPVTGVLDEETKEWMKRPRCGVPDQFGVRVKANLRRRRKRYALTGRKWNNHHL
MMP_14    67 AMQKFYGLQVTGKADADTMKAMRRPRCGVPDKFGAEIKAN--VRRKRYAIQGLKWQHNEI
MMP_16    75 AMQQFYGINMTGKVDRNTIDWMKKPRCGVPDQTRGSSKFH--IRRKRYALTGQKWQHKHI
MMP_24   113 TMQQFYGIPVTGVLDQTTIEWMKKPRCGVPDHPHLSR--R--RRNKRYALTGQKWRQKHI

MMP_15   145 TFSIQNYTEKLGWYHSMEAVRRAFRVWEQATPLVFQEVPYEDIRLRRQKEADIMVLFASG
MMP_14   125 TFCIQNYTPKVGEYATYEAIRKAFRVWESATPLRFREVPYAYIREGHEKQADIMIFFAEG
MMP_16   133 TYSIKNVTPKVGDPETRKAIRRAFDVWQNVTPLTFEEVPYSELENG-KRDVDITIIFASG
MMP_24   169 TYSIHNYTPKVGELDTRKAIRQAFDVWQKVTPLTFEEVPYHEIKSD-RKEADIMIFFASG

MMP_15   205 FHGDSSPFDGTGGFLAHAYFPGPGLGGDTHFDADEPWTFSSTDLHGNNLFLVAVHELGHA
MMP_14   185 FHGDSTPFDGEGGFLAHAYFPGPNIGGDTHFDSAEPWTVRNEDLNGNDIFLVAVHELGHA
MMP_16   192 FHGDSSPFDGEGGFLAHAYFPGPGIGGDTHFDSDEPWTLGNPNHDGNDLFLVAVHELGHA
MMP_24   228 FHGDSSPFDGEGGFLAHAYFPGPGIGGDTHFDSDEPWTLGNANHDGNDLFLVAVHELGHA

MMP_15   265 LGLEHSSNPNAIMAPFYQWKDVDNFKLPEDDLRGIQQLYGTPDGQPQPTQPLPTVTPRRP
MMP_14   245 LGLEHSSDPSAIMAPFYQWMDTENFVLPDDDRRGIQQLYGGESGFPTKMPPQPRT-----
MMP_16   252 LGLEHSNDPTAIMAPFYQYMETDNFKLPNDDLQGIQKIYGPPDKIPPPTRPLPTVPPHRS
MMP_24   288 LGLEHSSDPSAIMAPFYQYMETHNFKLPQDDLQGIQKIYGPPAEPLEPTRPLPTLPVRRI

MMP_15   325 GRPDHRPPRPPQPPPPGGKPERPPKPGPPVQPRATERPDQYGPNICDGDFDTVAMLRGEM
MMP_14   300 -------------------TSRPSVPDKP-------KNPTYGPNICDGNFDTVAMLRGEM
MMP_16   312 IPPADPRKNDR-P-----KPPRPPTG-RP-------SYPGAKPNICDGNFNTLAILRREM
MMP_24   348 HSPSERKHERQ-P-----RPPRPPLGDRP-------STPGTKPNICDGNFNTVALFRGEM

MMP_15   385 FVFKGRWFWRVRHNRVLDNYPMPIGHFWRGLPGDISAAYERQDGRFVFFKGDRYWLFREA
MMP_14   334 FVFKERWFWRVRNNQVMDGYPMPIGQFWRGLPASINTAYERKDGKFVFFKGDKHWVFDEA
MMP_16   358 FVFKDQWFWRVRNNRVMDGYPMQITYFWRGLPPSIDAVYENSDGNFVFFKGNKYWVFKDT
MMP_24   395 FVFKDRWFWRLRNNRVQEGYPMQIEQFWKGLPARIDAAYERADGRFVFFKGDKYWVFKEV

MMP_15   445 NLEPGYPQPLTSYGLGIPYDRIDTAIWWEPTGHTFFFQEDRYWRFNEETQRGDPGYPKPI
MMP_14   394 SLEPGYPKHIKELGRGLPTDKIDAALFWMPNGKTYFFRGNKYYRFNEELRAVDSEYPKNI
MMP_16   418 TLQPGYPHDLITLGSGIPPHGIDSAIWWEDVGKTYFFKGDRYWRYSEEMKTMDPGYPKPI
MMP_24   455 TVEPGYPHSLGELGSCLPREGIDTALRWEPVGKTYFFKGERYWRYSEERRATDPGYPKPI
```

```
MMP_15   505 SVWQGIPASPKGAFLSNDAAYTYFYKGTKYWKFDNERLRMEPGYPKSILRDFMGCQEHVE
MMP_14   454 KVWEGIPESPRGSFMGSDEVFTYFYKGNKYWKFNNQKLKVEPGYPKSALRDWMGCPSGG-
MMP_16   478 TVWKGIPESPQGAFVHKENGFTYFYKGKEYWKFNNQILKVEPGYPRSILKDFMGCDGPT-
MMP_24   515 TVWKGIPQAPQGAFISKEGYYTYFYKGRDYWKFDNQKLSVEPGYPRNILRDWMGCNQKEV

MMP_15   565 PGPRWPDVARPPFNPHGGAEPGADSAEGDVGDGDGDFGAGVNKDGGSRVVVQMEEVARTV
MMP_14   513 -RPD----------------------------------EGTEEETEVIIIEVDEEGGGAV
MMP_16   537 DRVK----------------------------------EGHSPPDDVDIVIKLDNTASTV
MMP_24   575 ERRK----------------------------------ERRLPQDDVDIMVTINDVPGSV

MMP_15   625 NVVMVLVPLLLLLCVLGLTYALVQMQRKGAPRVLLYCKRSLQEWV
MMP_14   538 SAAAVVLPVLLLLLVLAVGLAVFFFRRHGTPRRLLYCQRSLLDKV
MMP_16   563 KAIAIVIPCILALCLLVLVYTVFQFKRKGTPRHILYCKRSMQEWV
MMP_24   601 NAVAVVIPCILSLCILVLVYTIFQFKNKTGPQPVTYYKRPVQEWV
```

Supplemental Figure 6. Boxshade alignment of MT-MMP paralogs, labeled by color according to NCBI annotations. The key is the same as in the previous boxshade alignment of Supplemental Figure 3.

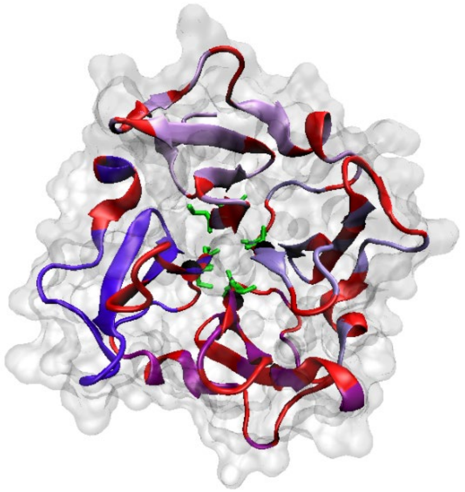

Supplemental Figure 7. The hemopexin domains of the MT-MMP paralog alignment are more variable than those of the mammalian ortholog alignment. The hemopexin domains of MMP-14 in surface and new cartoon representation. The metal binding sites are represented in green and variable regions of the MT-MMP alignment are represented in red.

#### b. Tables

Supplemental Table 1. Percent identity matrix of the mammalian MMP-15 ortholog alignment over the whole sequence, produced from Clustal Omega.

| | Mus musculus | Rattus norvegicus | Homo sapiens | Pan troglodytes | Canis lupus familiaris | Bos taurus | Equus caballus |
|---|---|---|---|---|---|---|---|
| Mus musculus | 100 | 97.72 | 89.38 | 89.54 | 88.8 | 89.33 | 88.26 |
| Rattus norvegicus | 97.72 | 100 | 88.92 | 89.08 | 88.48 | 89.33 | 88.57 |
| Homo sapiens | 89.38 | 88.92 | 100 | 99.55 | 92.42 | 94.29 | 92.83 |
| Pan troglodytes | 89.54 | 89.08 | 99.55 | 100 | 92.42 | 94.44 | 92.83 |
| Canis lupus familiaris | 88.8 | 88.48 | 92.42 | 92.42 | 100 | 94.84 | 93.15 |
| Bos taurus | 89.33 | 89.33 | 94.29 | 94.44 | 94.88 | 100 | 94.67 |
| Equus caballus | 88.26 | 88.57 | 92.83 | 92.83 | 93.15 | 94.67 | 100 |

Supplemental Table 2. Percent identity matrix of the mammalian MMP-15 ortholog alignment over the catalytic domain sequence, produced from Clustal Omega.

| | Mus musculus | Rattus norvegicus | Homo sapiens | Pan troglodytes | Bos taurus | Canis lupus familiaris | Equus caballus |
|---|---|---|---|---|---|---|---|
| Mus musculus | 100 | 98.8 | 90.96 | 90.96 | 92.77 | 92.77 | 92.77 |
| Rattus norvegicus | 98.8 | 100 | 90.96 | 90.96 | 92.77 | 92.77 | 93.98 |
| Homo sapiens | 90.96 | 90.96 | 100 | 99.4 | 96.41 | 95.81 | 94.61 |
| Pan troglodytes | 90.96 | 90.96 | 99.4 | 100 | 97.01 | 95.81 | 94.61 |
| Bos taurus | 92.77 | 92.77 | 96.41 | 97.01 | 100 | 98.8 | 97.6 |
| Canis lupus familiaris | 92.77 | 92.77 | 95.81 | 95.81 | 98.8 | 100 | 97.6 |
| Equus caballus | 92.77 | 93.98 | 94.61 | 94.61 | 97.6 | 97.6 | 100 |

Supplemental Table 3. Shannon Entropy values for the mammalian MMP-15 ortholog alignment, produced with the Protein Variability Server.

| Position | Consensus Sequence | Shannon Entropy |
|---|---|---|
| 1 | M | 0.592 |
| 2 | G | 0.592 |
| 3 | S | 0.592 |
| 4 | D | 0.592 |
| 5 | R | 1.149 |
| 6 | S | 0.592 |
| 7 | A | 0.592 |
| 8 | P | 1.449 |
| 9 | G | 1.149 |
| 10 | R | 0.592 |
| 11 | P | 0.592 |
| 12 | G | 0.592 |
| 13 | W | 1.842 |
| 14 | A | 1.449 |
| 15 | G | 0.592 |
| 16 | S | 0.592 |
| 17 | L | 1.842 |
| 18 | L | 0.592 |
| 19 | G | 1.379 |
| 20 | S | 1.95 |
| 21 | R | 0.985 |
| 22 | E | 0.985 |
| 23 | - | 1.557 |
| 24 | A | 0.985 |
| 25 | A | 1.449 |
| 26 | R | 0.592 |
| 27 | P | 1.379 |
| 28 | R | 1.842 |
| 29 | L | 1.149 |
| 30 | L | 0.592 |
| 31 | P | 1.149 |
| 32 | L | 0.592 |
| 33 | L | 0.592 |
| 34 | L | 0.592 |
| 35 | V | 0.592 |
| 36 | L | 0.592 |
| 37 | L | 0.592 |

| | | |
|---|---|---|
| 38 | G | 1.842 |
| 39 | C | 0.592 |
| 40 | L | 0.592 |
| 41 | G | 0 |
| 42 | R | 1.95 |
| 43 | G | 0 |
| 44 | A | 1.95 |
| 45 | A | 0.592 |
| 46 | A | 1.379 |
| 47 | E | 1.149 |
| 48 | D | 0.592 |
| 49 | A | 0.592 |
| 50 | E | 0.592 |
| 51 | V | 0 |
| 52 | - | 0.863 |
| 53 | H | 2.236 |
| 54 | A | 0 |
| 55 | E | 0.592 |
| 56 | N | 0 |
| 57 | W | 0 |
| 58 | L | 0 |
| 59 | R | 0 |
| 60 | L | 0 |
| 61 | Y | 0 |
| 62 | G | 0 |
| 63 | Y | 0 |
| 64 | L | 0 |
| 65 | P | 0 |
| 66 | Q | 0 |
| 67 | P | 0 |
| 68 | S | 0 |
| 69 | R | 0 |
| 70 | H | 0 |
| 71 | M | 0 |
| 72 | S | 0 |
| 73 | T | 0 |
| 74 | M | 0.592 |
| 75 | R | 0 |
| 76 | S | 0 |
| 77 | A | 0 |
| 78 | Q | 0 |
| 79 | I | 0.592 |
| 80 | L | 0 |

| | | |
|---|---|---|
| 81 | A | 0 |
| 82 | S | 0 |
| 83 | A | 0 |
| 84 | L | 0 |
| 85 | A | 0 |
| 86 | E | 0 |
| 87 | M | 0 |
| 88 | Q | 0 |
| 89 | R | 0.863 |
| 90 | F | 0 |
| 91 | Y | 0 |
| 92 | G | 0 |
| 93 | I | 0 |
| 94 | P | 0 |
| 95 | V | 0 |
| 96 | T | 0 |
| 97 | G | 0 |
| 98 | V | 0 |
| 99 | L | 0 |
| 100 | D | 0 |
| 101 | E | 0 |
| 102 | E | 0 |
| 103 | T | 0 |
| 104 | K | 0 |
| 105 | A | 1.557 |
| 106 | W | 0 |
| 107 | M | 0 |
| 108 | K | 0 |
| 109 | R | 0 |
| 110 | P | 0 |
| 111 | R | 0 |
| 112 | C | 0 |
| 113 | G | 0 |
| 114 | V | 0 |
| 115 | P | 0 |
| 116 | D | 0 |
| 117 | Q | 0 |
| 118 | F | 0 |
| 119 | G | 0 |
| 120 | V | 0 |
| 121 | R | 0.592 |
| 122 | V | 0 |
| 123 | K | 0 |

| | | |
|---|---|---|
| 124 | A | 0 |
| 125 | N | 0 |
| 126 | L | 0 |
| 127 | R | 0 |
| 128 | R | 0 |
| 129 | R | 0 |
| 130 | R | 0 |
| 131 | K | 0.863 |
| 132 | R | 0 |
| 133 | Y | 0 |
| 134 | A | 0.863 |
| 135 | L | 0 |
| 136 | T | 0 |
| 137 | G | 0 |
| 138 | R | 0.863 |
| 139 | K | 1.149 |
| 140 | W | 0 |
| 141 | N | 0.863 |
| 142 | N | 0.592 |
| 143 | H | 0.863 |
| 144 | H | 0 |
| 145 | L | 0 |
| 146 | T | 0 |
| 147 | F | 0 |
| 148 | S | 0 |
| 149 | I | 0 |
| 150 | Q | 0 |
| 151 | N | 0 |
| 152 | Y | 0 |
| 153 | T | 0 |
| 154 | E | 0 |
| 155 | K | 0 |
| 156 | L | 0 |
| 157 | G | 0 |
| 158 | W | 0 |
| 159 | Y | 0 |
| 160 | H | 0.863 |
| 161 | S | 0 |
| 162 | M | 0.985 |
| 163 | E | 0 |
| 164 | A | 0 |
| 165 | V | 0 |
| 166 | R | 0 |

| | | |
|---|---|---|
| 167 | R | 0 |
| 168 | A | 0 |
| 169 | F | 0 |
| 170 | R | 0.985 |
| 171 | V | 0 |
| 172 | W | 0 |
| 173 | E | 0 |
| 174 | Q | 0 |
| 175 | A | 0.863 |
| 176 | T | 0 |
| 177 | P | 0 |
| 178 | L | 0 |
| 179 | V | 0 |
| 180 | F | 0 |
| 181 | Q | 0 |
| 182 | E | 0 |
| 183 | V | 0 |
| 184 | P | 0.592 |
| 185 | Y | 0 |
| 186 | E | 0.863 |
| 187 | D | 0 |
| 188 | I | 0 |
| 189 | R | 0 |
| 190 | L | 0 |
| 191 | R | 0 |
| 192 | R | 0 |
| 193 | Q | 0.863 |
| 194 | K | 0.863 |
| 195 | E | 0 |
| 196 | A | 0 |
| 197 | D | 0 |
| 198 | I | 0 |
| 199 | M | 0 |
| 200 | V | 0 |
| 201 | L | 0 |
| 202 | F | 0 |
| 203 | A | 0 |
| 204 | S | 0 |
| 205 | G | 0 |
| 206 | F | 0 |
| 207 | H | 0 |
| 208 | G | 0 |
| 209 | D | 0 |

| | | |
|---|---|---|
| 210 | S | 0 |
| 211 | S | 0 |
| 212 | P | 0 |
| 213 | F | 0 |
| 214 | D | 0 |
| 215 | G | 0 |
| 216 | T | 0.863 |
| 217 | G | 0 |
| 218 | G | 0 |
| 219 | F | 0 |
| 220 | L | 0 |
| 221 | A | 0 |
| 222 | H | 0 |
| 223 | A | 0 |
| 224 | Y | 0 |
| 225 | F | 0 |
| 226 | P | 0 |
| 227 | G | 0 |
| 228 | P | 0 |
| 229 | G | 0 |
| 230 | L | 0 |
| 231 | G | 0 |
| 232 | G | 0 |
| 233 | D | 0 |
| 234 | T | 0 |
| 235 | H | 0 |
| 236 | F | 0 |
| 237 | D | 0 |
| 238 | A | 0 |
| 239 | D | 0 |
| 240 | E | 0 |
| 241 | P | 0 |
| 242 | W | 0 |
| 243 | T | 0 |
| 244 | F | 0 |
| 245 | S | 0 |
| 246 | S | 0 |
| 247 | T | 0 |
| 248 | D | 0 |
| 249 | L | 0 |
| 250 | H | 0 |
| 251 | G | 0 |
| 252 | N | 0.863 |

| | | |
|---|---|---|
| 253 | S | 0.863 |
| 254 | L | 0 |
| 255 | F | 0 |
| 256 | L | 0 |
| 257 | V | 0 |
| 258 | A | 0 |
| 259 | V | 0 |
| 260 | H | 0 |
| 261 | E | 0 |
| 262 | L | 0 |
| 263 | G | 0 |
| 264 | H | 0 |
| 265 | A | 0 |
| 266 | L | 0 |
| 267 | G | 0 |
| 268 | L | 0 |
| 269 | E | 0 |
| 270 | H | 0 |
| 271 | S | 0 |
| 272 | S | 0 |
| 273 | N | 0 |
| 274 | P | 0 |
| 275 | S | 0.863 |
| 276 | A | 0 |
| 277 | I | 0 |
| 278 | M | 0 |
| 279 | A | 0 |
| 280 | P | 0 |
| 281 | F | 0 |
| 282 | Y | 0 |
| 283 | Q | 0 |
| 284 | W | 0 |
| 285 | M | 0.863 |
| 286 | D | 0 |
| 287 | T | 1.379 |
| 288 | D | 0 |
| 289 | N | 0.592 |
| 290 | F | 0 |
| 291 | Q | 0.863 |
| 292 | L | 0 |
| 293 | P | 0 |
| 294 | E | 0 |
| 295 | D | 0 |

| | | |
|---|---|---|
| 296 | D | 0 |
| 297 | L | 0 |
| 298 | R | 0 |
| 299 | G | 0 |
| 300 | I | 0 |
| 301 | Q | 0 |
| 302 | Q | 0 |
| 303 | L | 0 |
| 304 | Y | 0 |
| 305 | G | 0 |
| 306 | T | 0.863 |
| 307 | P | 0 |
| 308 | D | 0 |
| 309 | G | 0 |
| 310 | Q | 0.863 |
| 311 | P | 0 |
| 312 | Q | 0 |
| 313 | P | 0 |
| 314 | T | 0 |
| 315 | R | 0.985 |
| 316 | P | 0.592 |
| 317 | L | 0 |
| 318 | P | 0 |
| 319 | T | 0 |
| 320 | V | 0 |
| 321 | T | 0.863 |
| 322 | P | 0 |
| 323 | R | 0 |
| 324 | R | 0 |
| 325 | P | 0 |
| 326 | G | 0 |
| 327 | R | 0 |
| 328 | P | 0 |
| 329 | D | 0 |
| 330 | H | 0 |
| 331 | R | 0.863 |
| 332 | P | 0 |
| 333 | P | 0 |
| 334 | R | 0 |
| 335 | P | 0 |
| 336 | P | 0 |
| 337 | Q | 0 |
| 338 | P | 0 |

| | | |
|---|---|---|
| 339 | P | 0 |
| 340 | P | 0.863 |
| 341 | P | 0 |
| 342 | G | 0 |
| 343 | G | 0 |
| 344 | K | 0 |
| 345 | P | 0 |
| 346 | E | 0 |
| 347 | R | 0 |
| 348 | P | 0 |
| 349 | P | 0 |
| 350 | K | 0 |
| 351 | P | 0 |
| 352 | G | 0 |
| 353 | P | 0 |
| 354 | P | 0 |
| 355 | P | 1.557 |
| 356 | Q | 0 |
| 357 | P | 0 |
| 358 | R | 0 |
| 359 | A | 0 |
| 360 | T | 0 |
| 361 | E | 0 |
| 362 | R | 0 |
| 363 | P | 0 |
| 364 | D | 0 |
| 365 | Q | 0 |
| 366 | Y | 0 |
| 367 | G | 0 |
| 368 | P | 0 |
| 369 | N | 0 |
| 370 | I | 0 |
| 371 | C | 0 |
| 372 | D | 0 |
| 373 | G | 0 |
| 374 | D | 0.863 |
| 375 | F | 0 |
| 376 | D | 0 |
| 377 | T | 0 |
| 378 | V | 0 |
| 379 | A | 0 |
| 380 | M | 0.863 |
| 381 | L | 0 |

| | | |
|---|---|---|
| 382 | R | 0 |
| 383 | G | 0 |
| 384 | E | 0 |
| 385 | M | 0 |
| 386 | F | 0 |
| 387 | V | 0 |
| 388 | F | 0 |
| 389 | K | 0 |
| 390 | G | 0 |
| 391 | R | 0 |
| 392 | W | 0 |
| 393 | F | 0 |
| 394 | W | 0 |
| 395 | R | 0 |
| 396 | V | 0 |
| 397 | R | 0 |
| 398 | H | 0 |
| 399 | N | 0 |
| 400 | R | 0 |
| 401 | V | 0 |
| 402 | L | 0 |
| 403 | D | 0 |
| 404 | N | 0.592 |
| 405 | Y | 0 |
| 406 | P | 0 |
| 407 | M | 0 |
| 408 | P | 0 |
| 409 | I | 0 |
| 410 | G | 0 |
| 411 | H | 0 |
| 412 | F | 0 |
| 413 | W | 0 |
| 414 | R | 0 |
| 415 | G | 0 |
| 416 | L | 0 |
| 417 | P | 0 |
| 418 | G | 0.863 |
| 419 | D | 0.863 |
| 420 | I | 0 |
| 421 | S | 0 |
| 422 | A | 0 |
| 423 | A | 0 |
| 424 | Y | 0 |

| | | |
|---|---|---|
| 425 | E | 0 |
| 426 | R | 0 |
| 427 | Q | 0 |
| 428 | D | 0 |
| 429 | G | 0 |
| 430 | R | 0.863 |
| 431 | F | 0 |
| 432 | V | 0 |
| 433 | F | 0 |
| 434 | F | 0 |
| 435 | K | 0 |
| 436 | G | 0 |
| 437 | D | 0.863 |
| 438 | R | 0 |
| 439 | Y | 0 |
| 440 | W | 0 |
| 441 | L | 0 |
| 442 | F | 0 |
| 443 | R | 0 |
| 444 | E | 0 |
| 445 | A | 0 |
| 446 | N | 0 |
| 447 | L | 0 |
| 448 | E | 0 |
| 449 | P | 0 |
| 450 | G | 0 |
| 451 | Y | 0 |
| 452 | P | 0 |
| 453 | Q | 0 |
| 454 | P | 0 |
| 455 | L | 0 |
| 456 | T | 0.863 |
| 457 | S | 0 |
| 458 | Y | 0 |
| 459 | G | 0 |
| 460 | L | 0.863 |
| 461 | G | 0.863 |
| 462 | I | 0 |
| 463 | P | 0 |
| 464 | Y | 0 |
| 465 | D | 0 |
| 466 | R | 0.592 |
| 467 | I | 0 |

| | | |
|---|---|---|
| 468 | D | 0 |
| 469 | T | 0 |
| 470 | A | 0 |
| 471 | I | 0 |
| 472 | W | 0 |
| 473 | W | 0 |
| 474 | E | 0 |
| 475 | P | 0 |
| 476 | T | 0 |
| 477 | G | 0 |
| 478 | H | 0 |
| 479 | T | 0 |
| 480 | F | 0 |
| 481 | F | 0 |
| 482 | F | 0 |
| 483 | Q | 0 |
| 484 | E | 0.863 |
| 485 | D | 0 |
| 486 | R | 0 |
| 487 | Y | 0 |
| 488 | W | 0 |
| 489 | R | 0 |
| 490 | F | 0 |
| 491 | N | 0 |
| 492 | E | 0 |
| 493 | E | 0 |
| 494 | T | 0 |
| 495 | Q | 0 |
| 496 | R | 0.863 |
| 497 | G | 0 |
| 498 | D | 0 |
| 499 | P | 0 |
| 500 | G | 0 |
| 501 | Y | 0 |
| 502 | P | 0 |
| 503 | K | 0 |
| 504 | P | 0 |
| 505 | I | 0 |
| 506 | S | 0 |
| 507 | V | 0 |
| 508 | W | 0 |
| 509 | Q | 0 |
| 510 | G | 0 |

| | | |
|---|---|---|
| 511 | I | 0 |
| 512 | P | 0 |
| 513 | A | 1.449 |
| 514 | S | 0 |
| 515 | P | 0 |
| 516 | K | 0 |
| 517 | G | 0 |
| 518 | A | 0 |
| 519 | F | 0 |
| 520 | L | 0 |
| 521 | S | 0 |
| 522 | N | 0 |
| 523 | D | 0 |
| 524 | A | 0 |
| 525 | A | 0 |
| 526 | Y | 0 |
| 527 | T | 0 |
| 528 | Y | 0 |
| 529 | F | 0 |
| 530 | Y | 0 |
| 531 | K | 0 |
| 532 | G | 0 |
| 533 | T | 0 |
| 534 | K | 0 |
| 535 | Y | 0 |
| 536 | W | 0 |
| 537 | K | 0 |
| 538 | F | 0 |
| 539 | D | 0.863 |
| 540 | N | 0 |
| 541 | E | 0 |
| 542 | R | 0 |
| 543 | L | 0 |
| 544 | R | 0 |
| 545 | M | 0 |
| 546 | E | 0 |
| 547 | P | 0 |
| 548 | G | 0 |
| 549 | Y | 0.863 |
| 550 | P | 0 |
| 551 | K | 0 |
| 552 | S | 0 |
| 553 | I | 0 |

| | | |
|---|---|---|
| 554 | L | 0 |
| 555 | R | 0 |
| 556 | D | 0 |
| 557 | F | 0 |
| 558 | M | 0 |
| 559 | G | 0 |
| 560 | C | 0 |
| 561 | Q | 0 |
| 562 | E | 0 |
| 563 | H | 0.592 |
| 564 | V | 0 |
| 565 | E | 0 |
| 566 | P | 0 |
| 567 | G | 0.863 |
| 568 | P | 0.863 |
| 569 | R | 0 |
| 570 | W | 0 |
| 571 | P | 0 |
| 572 | D | 0 |
| 573 | V | 0 |
| 574 | A | 0 |
| 575 | R | 0 |
| 576 | P | 0 |
| 577 | P | 0 |
| 578 | F | 0 |
| 579 | N | 0 |
| 580 | P | 0 |
| 581 | D | 1.557 |
| 582 | G | 0 |
| 583 | G | 0 |
| 584 | A | 0 |
| 585 | E | 0 |
| 586 | P | 0 |
| 587 | G | 0.863 |
| 588 | A | 1.149 |
| 589 | D | 1.449 |
| 590 | G | 1.379 |
| 591 | D | 0.863 |
| 592 | S | 1.379 |
| 593 | E | 1.557 |
| 594 | E | 0.863 |
| 595 | G | 1.557 |
| 596 | - | 2.236 |

| | | |
|---|---|---|
| 597 | E | 1.842 |
| 598 | - | 0.985 |
| 599 | - | 1.665 |
| 600 | E | 1.379 |
| 601 | A | 1.95 |
| 602 | G | 1.557 |
| 603 | - | 1.95 |
| 604 | G | 0.863 |
| 605 | G | 1.557 |
| 606 | G | 1.379 |
| 607 | D | 1.842 |
| 608 | G | 1.842 |
| 609 | D | 0.985 |
| 610 | F | 1.449 |
| 611 | G | 0.985 |
| 612 | A | 1.842 |
| 613 | G | 1.379 |
| 614 | T | 1.95 |
| 615 | D | 0.863 |
| 616 | K | 1.379 |
| 617 | D | 0 |
| 618 | G | 0.863 |
| 619 | G | 0 |
| 620 | S | 0 |
| 621 | R | 0 |
| 622 | V | 0 |
| 623 | V | 0 |
| 624 | V | 0 |
| 625 | Q | 0 |
| 626 | M | 0.592 |
| 627 | E | 0 |
| 628 | E | 0 |
| 629 | V | 0 |
| 630 | T | 1.557 |
| 631 | R | 0 |
| 632 | T | 0 |
| 633 | V | 0 |
| 634 | N | 0.592 |
| 635 | V | 0.985 |
| 636 | V | 0 |
| 637 | M | 0 |
| 638 | V | 0 |
| 639 | L | 0 |

| | | |
|---|---|---|
| 640 | V | 0 |
| 641 | P | 0 |
| 642 | - | 0.985 |
| 643 | L | 0.592 |
| 644 | L | 0 |
| 645 | L | 0 |
| 646 | L | 0 |
| 647 | L | 0 |
| 648 | C | 0 |
| 649 | I | 0.985 |
| 650 | L | 0 |
| 651 | G | 0 |
| 652 | L | 0 |
| 653 | T | 0.863 |
| 654 | Y | 0.863 |
| 655 | A | 0 |
| 656 | L | 0 |
| 657 | V | 0 |
| 658 | Q | 0.592 |
| 659 | M | 0 |
| 660 | Q | 0 |
| 661 | R | 0 |
| 662 | K | 0 |
| 663 | G | 0 |
| 664 | A | 0 |
| 665 | P | 0 |
| 666 | R | 0 |
| 667 | M | 0.985 |
| 668 | L | 0 |
| 669 | L | 0 |
| 670 | Y | 0 |
| 671 | C | 0 |
| 672 | K | 0 |
| 673 | R | 0 |
| 674 | S | 0 |
| 675 | L | 0 |
| 676 | Q | 0 |
| 677 | E | 0 |
| 678 | W | 0 |
| 679 | V | 0 |

Supplemental Table 4. Percent similarity matrix of the MT-MMP paralog alignment over the entire sequence, produced using Clustal Omega.

| | MMP_15 | MMP_14 | MMP_16 | MMP_24 |
|---|---|---|---|---|
| MMP_15 | 100 | 58.33 | 56.51 | 55.74 |
| MMP_14 | 58.33 | 100 | 56.24 | 55.79 |
| MMP_16 | 56.51 | 56.24 | 100 | 66.39 |
| MMP_24 | 55.74 | 55.79 | 66.39 | 100 |

Supplemental Table 5. Shannon Entropy values for the MT-MMP paralog alignment, produced using the Protein Variability Server.

| Position | MMP-15 Sequence | Shannon Entropy |
|---|---|---|
| 1 | - | 0.811 |
| 2 | - | 0.811 |
| 3 | - | 0.811 |
| 4 | - | 0.811 |
| 5 | - | 0.811 |
| 6 | - | 0.811 |
| 7 | - | 0.811 |
| 8 | - | 0.811 |
| 9 | M | 1.5 |
| 10 | G | 2 |
| 11 | S | 1.5 |
| 12 | D | 2 |
| 13 | P | 1 |
| 14 | - | 0.811 |
| 15 | - | 0.811 |
| 16 | - | 0.811 |
| 17 | - | 1 |
| 18 | S | 1.5 |
| 19 | A | 1.5 |
| 20 | P | 1 |
| 21 | - | 0.811 |
| 22 | G | 1.5 |
| 23 | R | 1.5 |
| 24 | P | 1 |
| 25 | G | 1.5 |
| 26 | W | 1 |
| 27 | T | 1.5 |
| 28 | G | 1.5 |
| 29 | S | 1.5 |

| | | |
|---|---|---|
| 30 | L | 0.811 |
| 31 | L | 0.811 |
| 32 | G | 0.811 |
| 33 | D | 0.811 |
| 34 | R | 0.811 |
| 35 | E | 0.811 |
| 36 | E | 0.811 |
| 37 | A | 1.5 |
| 38 | A | 2 |
| 39 | R | 1.5 |
| 40 | P | 1.5 |
| 41 | R | 0.811 |
| 42 | L | 1.5 |
| 43 | L | 0.811 |
| 44 | P | 0.811 |
| 45 | L | 0 |
| 46 | L | 1.5 |
| 47 | L | 0.811 |
| 48 | V | 2 |
| 49 | L | 2 |
| 50 | L | 0.811 |
| 51 | G | 1.5 |
| 52 | C | 1 |
| 53 | L | 1 |
| 54 | G | 2 |
| 55 | - | 0.811 |
| 56 | - | 1.5 |
| 57 | - | 1.5 |
| 58 | - | 1.5 |
| 59 | - | 1.5 |
| 60 | - | 1.5 |
| 61 | - | 1.5 |
| 62 | - | 1.5 |
| 63 | - | 1.5 |
| 64 | - | 1.5 |
| 65 | - | 1 |
| 66 | - | 1.5 |
| 67 | - | 1.5 |
| 68 | - | 0.811 |
| 69 | - | 1.5 |
| 70 | - | 1.5 |
| 71 | - | 1.5 |
| 72 | - | 1.5 |

| | | |
|---|---|---|
| 73 | - | 1.5 |
| 74 | - | 2 |
| 75 | - | 2 |
| 76 | - | 1.5 |
| 77 | - | 2 |
| 78 | L | 1.5 |
| 79 | G | 2 |
| 80 | V | 1.5 |
| 81 | A | 2 |
| 82 | A | 1.5 |
| 83 | E | 2 |
| 84 | D | 1.5 |
| 85 | A | 1.5 |
| 86 | E | 2 |
| 87 | V | 0.811 |
| 88 | H | 2 |
| 89 | A | 2 |
| 90 | E | 0.811 |
| 91 | N | 1.5 |
| 92 | W | 0 |
| 93 | L | 0 |
| 94 | R | 1.5 |
| 95 | L | 2 |
| 96 | Y | 0 |
| 97 | G | 0 |
| 98 | Y | 0 |
| 99 | L | 0 |
| 100 | P | 0.811 |
| 101 | Q | 0.811 |
| 102 | P | 2 |
| 103 | S | 0.811 |
| 104 | R | 2 |
| 105 | H | 0.811 |
| 106 | M | 1.5 |
| 107 | S | 0.811 |
| 108 | T | 1.5 |
| 109 | M | 1.5 |
| 110 | R | 0.811 |
| 111 | S | 0 |
| 112 | A | 0.811 |
| 113 | Q | 1.5 |
| 114 | I | 2 |
| 115 | L | 0.811 |

| | | |
|---|---|---|
| 116 | A | 1.5 |
| 117 | S | 0.811 |
| 118 | A | 0 |
| 119 | L | 1.5 |
| 120 | A | 0.811 |
| 121 | E | 1.5 |
| 122 | M | 0 |
| 123 | Q | 0 |
| 124 | R | 1.5 |
| 125 | F | 0 |
| 126 | Y | 0 |
| 127 | G | 0 |
| 128 | I | 0.811 |
| 129 | P | 1.5 |
| 130 | V | 0.811 |
| 131 | T | 0 |
| 132 | G | 0 |
| 133 | V | 1 |
| 134 | L | 1.5 |
| 135 | D | 0 |
| 136 | E | 2 |
| 137 | E | 2 |
| 138 | T | 0 |
| 139 | K | 1.5 |
| 140 | E | 1.5 |
| 141 | W | 0.811 |
| 142 | M | 0 |
| 143 | K | 0.811 |
| 144 | R | 1 |
| 145 | P | 0 |
| 146 | R | 0 |
| 147 | C | 0 |
| 148 | G | 0 |
| 149 | V | 0 |
| 150 | P | 0 |
| 151 | D | 0 |
| 152 | Q | 1.5 |
| 153 | F | 1.5 |
| 154 | G | 1.5 |
| 155 | V | 2 |
| 156 | R | 1.5 |
| 157 | V | 2 |
| 158 | K | 0.811 |

| | | |
|---|---|---|
| 159 | A | 1.5 |
| 160 | N | 1.5 |
| 161 | L | 0.811 |
| 162 | R | 0.811 |
| 163 | R | 1.5 |
| 164 | R | 0 |
| 165 | R | 0.811 |
| 166 | K | 0 |
| 167 | R | 0 |
| 168 | Y | 0 |
| 169 | A | 0 |
| 170 | L | 0.811 |
| 171 | T | 0.811 |
| 172 | G | 0 |
| 173 | R | 1.5 |
| 174 | K | 0 |
| 175 | W | 0 |
| 176 | N | 1.5 |
| 177 | N | 1.5 |
| 178 | H | 1.5 |
| 179 | H | 0.811 |
| 180 | L | 0.811 |
| 181 | T | 0 |
| 182 | F | 1 |
| 183 | S | 0.811 |
| 184 | I | 0 |
| 185 | Q | 1.5 |
| 186 | N | 0 |
| 187 | Y | 0.811 |
| 188 | T | 0 |
| 189 | E | 0.811 |
| 190 | K | 0 |
| 191 | L | 0.811 |
| 192 | G | 0 |
| 193 | W | 1.5 |
| 194 | Y | 1.5 |
| 195 | H | 2 |
| 196 | S | 0.811 |
| 197 | M | 1.5 |
| 198 | E | 1 |
| 199 | A | 0 |
| 200 | V | 0.811 |
| 201 | R | 0 |

| | | |
|---|---|---|
| 202 | R | 1.5 |
| 203 | A | 0 |
| 204 | F | 0 |
| 205 | R | 1 |
| 206 | V | 0 |
| 207 | W | 0 |
| 208 | E | 1 |
| 209 | Q | 2 |
| 210 | A | 1 |
| 211 | T | 0 |
| 212 | P | 0 |
| 213 | L | 0 |
| 214 | V | 1.5 |
| 215 | F | 0 |
| 216 | Q | 1.5 |
| 217 | E | 0 |
| 218 | V | 0 |
| 219 | P | 0 |
| 220 | Y | 0 |
| 221 | E | 2 |
| 222 | D | 1.5 |
| 223 | I | 0.811 |
| 224 | R | 1.5 |
| 225 | L | 2 |
| 226 | R | 1.5 |
| 227 | R | 1.5 |
| 228 | Q | 2 |
| 229 | K | 0.811 |
| 230 | E | 1.5 |
| 231 | A | 0.811 |
| 232 | D | 0 |
| 233 | I | 0 |
| 234 | M | 0.811 |
| 235 | V | 0.811 |
| 236 | L | 1.5 |
| 237 | F | 0 |
| 238 | A | 0 |
| 239 | S | 0.811 |
| 240 | G | 0 |
| 241 | F | 0 |
| 242 | H | 0 |
| 243 | G | 0 |
| 244 | D | 0 |

| | | |
|---|---|---|
| 245 | S | 0 |
| 246 | S | 0.811 |
| 247 | P | 0 |
| 248 | F | 0 |
| 249 | D | 0 |
| 250 | G | 0 |
| 251 | T | 0.811 |
| 252 | G | 0 |
| 253 | G | 0 |
| 254 | F | 0 |
| 255 | L | 0 |
| 256 | A | 0 |
| 257 | H | 0 |
| 258 | A | 0 |
| 259 | Y | 0 |
| 260 | F | 0 |
| 261 | P | 0 |
| 262 | G | 0 |
| 263 | P | 0 |
| 264 | G | 0.811 |
| 265 | L | 0.811 |
| 266 | G | 0 |
| 267 | G | 0 |
| 268 | D | 0 |
| 269 | T | 0 |
| 270 | H | 0 |
| 271 | F | 0 |
| 272 | D | 0 |
| 273 | A | 0.811 |
| 274 | D | 0.811 |
| 275 | E | 0 |
| 276 | P | 0 |
| 277 | W | 0 |
| 278 | T | 0 |
| 279 | F | 1.5 |
| 280 | S | 1.5 |
| 281 | S | 0.811 |
| 282 | T | 2 |
| 283 | D | 1 |
| 284 | L | 1 |
| 285 | H | 1.5 |
| 286 | G | 0 |
| 287 | N | 0 |

| | | |
|---|---|---|
| 288 | N | 0.811 |
| 289 | L | 0.811 |
| 290 | F | 0 |
| 291 | L | 0 |
| 292 | V | 0 |
| 293 | A | 0 |
| 294 | V | 0 |
| 295 | H | 0 |
| 296 | E | 0 |
| 297 | L | 0 |
| 298 | G | 0 |
| 299 | H | 0 |
| 300 | A | 0 |
| 301 | L | 0 |
| 302 | G | 0 |
| 303 | L | 0 |
| 304 | E | 0 |
| 305 | H | 0 |
| 306 | S | 0 |
| 307 | S | 0.811 |
| 308 | N | 0.811 |
| 309 | P | 0 |
| 310 | N | 1.5 |
| 311 | A | 0 |
| 312 | I | 0 |
| 313 | M | 0 |
| 314 | A | 0 |
| 315 | P | 0 |
| 316 | F | 0 |
| 317 | Y | 0 |
| 318 | Q | 0 |
| 319 | W | 1 |
| 320 | K | 0.811 |
| 321 | D | 1 |
| 322 | V | 0.811 |
| 323 | D | 1.5 |
| 324 | N | 0 |
| 325 | F | 0 |
| 326 | K | 0.811 |
| 327 | L | 0 |
| 328 | P | 0 |
| 329 | E | 2 |
| 330 | D | 0 |

| | | |
|---|---|---|
| 331 | D | 0 |
| 332 | L | 0.811 |
| 333 | R | 1 |
| 334 | G | 0 |
| 335 | I | 0 |
| 336 | Q | 0 |
| 337 | Q | 1 |
| 338 | L | 1 |
| 339 | Y | 0 |
| 340 | G | 0 |
| 341 | T | 1.5 |
| 342 | P | 0.811 |
| 343 | D | 1.5 |
| 344 | G | 1.5 |
| 345 | Q | 2 |
| 346 | P | 0.811 |
| 347 | Q | 2 |
| 348 | P | 0.811 |
| 349 | T | 0.811 |
| 350 | Q | 1.5 |
| 351 | P | 0 |
| 352 | L | 0.811 |
| 353 | P | 0 |
| 354 | T | 0.811 |
| 355 | V | 1.5 |
| 356 | T | 1.5 |
| 357 | P | 1.5 |
| 358 | R | 1.5 |
| 359 | R | 0.811 |
| 360 | P | 2 |
| 361 | G | 2 |
| 362 | R | 2 |
| 363 | P | 0.811 |
| 364 | D | 2 |
| 365 | H | 2 |
| 366 | R | 1.5 |
| 367 | P | 2 |
| 368 | P | 2 |
| 369 | R | 2 |
| 370 | P | 2 |
| 371 | P | 2 |
| 372 | Q | 0.811 |
| 373 | P | 0.811 |

| | | |
|---|---|---|
| 374 | P | 0.811 |
| 375 | P | 0.811 |
| 376 | P | 0.811 |
| 377 | G | 0.811 |
| 378 | G | 0.811 |
| 379 | K | 1.5 |
| 380 | P | 0.811 |
| 381 | E | 1.5 |
| 382 | R | 0 |
| 383 | P | 0 |
| 384 | P | 0.811 |
| 385 | K | 2 |
| 386 | P | 1 |
| 387 | G | 1.5 |
| 388 | P | 1.5 |
| 389 | P | 0 |
| 390 | V | 0.811 |
| 391 | Q | 0.811 |
| 392 | P | 0.811 |
| 393 | R | 0.811 |
| 394 | A | 0.811 |
| 395 | T | 0.811 |
| 396 | E | 0.811 |
| 397 | R | 1.5 |
| 398 | P | 2 |
| 399 | D | 0.811 |
| 400 | Q | 1.5 |
| 401 | Y | 1.5 |
| 402 | G | 1 |
| 403 | P | 0 |
| 404 | N | 0 |
| 405 | I | 0 |
| 406 | C | 0 |
| 407 | D | 0 |
| 408 | G | 0 |
| 409 | D | 0.811 |
| 410 | F | 0 |
| 411 | D | 1 |
| 412 | T | 0 |
| 413 | V | 0.811 |
| 414 | A | 0 |
| 415 | M | 1.5 |
| 416 | L | 0.811 |

| | | |
|---|---|---|
| 417 | R | 0 |
| 418 | G | 0.811 |
| 419 | E | 0 |
| 420 | M | 0 |
| 421 | F | 0 |
| 422 | V | 0 |
| 423 | F | 0 |
| 424 | K | 0 |
| 425 | G | 1.5 |
| 426 | R | 0.811 |
| 427 | W | 0 |
| 428 | F | 0 |
| 429 | W | 0 |
| 430 | R | 0 |
| 431 | V | 0.811 |
| 432 | R | 0 |
| 433 | H | 0.811 |
| 434 | N | 0 |
| 435 | R | 0.811 |
| 436 | V | 0 |
| 437 | L | 1.5 |
| 438 | D | 0.811 |
| 439 | N | 0.811 |
| 440 | Y | 0 |
| 441 | P | 0 |
| 442 | M | 0 |
| 443 | P | 1 |
| 444 | I | 0 |
| 445 | G | 1.5 |
| 446 | H | 1.5 |
| 447 | F | 0 |
| 448 | W | 0 |
| 449 | R | 0.811 |
| 450 | G | 0 |
| 451 | L | 0 |
| 452 | P | 0 |
| 453 | G | 1.5 |
| 454 | D | 1.5 |
| 455 | I | 0 |
| 456 | S | 1.5 |
| 457 | A | 0.811 |
| 458 | A | 0.811 |
| 459 | Y | 0 |

| | | |
|---|---|---|
| 460 | E | 0 |
| 461 | R | 0.811 |
| 462 | Q | 2 |
| 463 | D | 0 |
| 464 | G | 0 |
| 465 | R | 1.5 |
| 466 | F | 0 |
| 467 | V | 0 |
| 468 | F | 0 |
| 469 | F | 0 |
| 470 | K | 0 |
| 471 | G | 0 |
| 472 | D | 0.811 |
| 473 | R | 0.811 |
| 474 | Y | 0.811 |
| 475 | W | 0 |
| 476 | L | 0.811 |
| 477 | F | 0 |
| 478 | R | 1.5 |
| 479 | E | 0.811 |
| 480 | A | 1.5 |
| 481 | N | 1.5 |
| 482 | L | 0.811 |
| 483 | E | 0.811 |
| 484 | P | 0 |
| 485 | G | 0 |
| 486 | Y | 0 |
| 487 | P | 0 |
| 488 | Q | 1.5 |
| 489 | P | 2 |
| 490 | L | 0.811 |
| 491 | T | 2 |
| 492 | S | 1.5 |
| 493 | Y | 0.811 |
| 494 | G | 0 |
| 495 | L | 1.5 |
| 496 | G | 0.811 |
| 497 | I | 1 |
| 498 | P | 0 |
| 499 | Y | 2 |
| 500 | D | 1.5 |
| 501 | R | 1.5 |
| 502 | I | 0 |

| | | |
|---|---|---|
| 503 | D | 0 |
| 504 | T | 1.5 |
| 505 | A | 0 |
| 506 | I | 1 |
| 507 | W | 1.5 |
| 508 | W | 0 |
| 509 | E | 0.811 |
| 510 | P | 0.811 |
| 511 | T | 1.5 |
| 512 | G | 0 |
| 513 | H | 0.811 |
| 514 | T | 0 |
| 515 | F | 0.811 |
| 516 | F | 0 |
| 517 | F | 0 |
| 518 | Q | 1.5 |
| 519 | E | 0.811 |
| 520 | D | 1.5 |
| 521 | R | 0.811 |
| 522 | Y | 0 |
| 523 | W | 0.811 |
| 524 | R | 0 |
| 525 | F | 1 |
| 526 | N | 1 |
| 527 | E | 0 |
| 528 | E | 0 |
| 529 | T | 2 |
| 530 | Q | 1.5 |
| 531 | R | 1.5 |
| 532 | G | 2 |
| 533 | D | 0 |
| 534 | P | 0.811 |
| 535 | G | 0.811 |
| 536 | Y | 0 |
| 537 | P | 0 |
| 538 | K | 0 |
| 539 | P | 0.811 |
| 540 | I | 0 |
| 541 | S | 1.5 |
| 542 | V | 0 |
| 543 | W | 0 |
| 544 | Q | 1.5 |
| 545 | G | 0 |

| | | |
|---|---|---|
| 546 | I | 0 |
| 547 | P | 0 |
| 548 | A | 1.5 |
| 549 | S | 0.811 |
| 550 | P | 0 |
| 551 | K | 1.5 |
| 552 | G | 0 |
| 553 | A | 0.811 |
| 554 | F | 0 |
| 555 | L | 2 |
| 556 | S | 1.5 |
| 557 | N | 1.5 |
| 558 | D | 1 |
| 559 | A | 2 |
| 560 | A | 2 |
| 561 | Y | 1 |
| 562 | T | 0 |
| 563 | Y | 0 |
| 564 | F | 0 |
| 565 | Y | 0 |
| 566 | K | 0 |
| 567 | G | 0 |
| 568 | T | 2 |
| 569 | K | 1.5 |
| 570 | Y | 0 |
| 571 | W | 0 |
| 572 | K | 0 |
| 573 | F | 0 |
| 574 | D | 1 |
| 575 | N | 0 |
| 576 | E | 0.811 |
| 577 | R | 1.5 |
| 578 | L | 0 |
| 579 | R | 1.5 |
| 580 | M | 0.811 |
| 581 | E | 0 |
| 582 | P | 0 |
| 583 | G | 0 |
| 584 | Y | 0 |
| 585 | P | 0 |
| 586 | K | 1 |
| 587 | S | 0.811 |
| 588 | I | 0.811 |

| | | |
|---|---|---|
| 589 | L | 0 |
| 590 | R | 0.811 |
| 591 | D | 0 |
| 592 | F | 1 |
| 593 | M | 0 |
| 594 | G | 0 |
| 595 | C | 0 |
| 596 | Q | 2 |
| 597 | E | 2 |
| 598 | H | 2 |
| 599 | V | 2 |
| 600 | E | 1.5 |
| 601 | P | 2 |
| 602 | G | 0.811 |
| 603 | P | 1.5 |
| 604 | R | 1.5 |
| 605 | W | 0.811 |
| 606 | P | 0.811 |
| 607 | D | 0.811 |
| 608 | V | 0.811 |
| 609 | A | 0.811 |
| 610 | R | 0.811 |
| 611 | P | 0.811 |
| 612 | P | 0.811 |
| 613 | F | 0.811 |
| 614 | N | 0.811 |
| 615 | P | 0.811 |
| 616 | H | 0.811 |
| 617 | G | 0.811 |
| 618 | G | 0.811 |
| 619 | A | 0.811 |
| 620 | E | 0.811 |
| 621 | P | 0.811 |
| 622 | G | 0.811 |
| 623 | A | 0.811 |
| 624 | D | 0.811 |
| 625 | S | 0.811 |
| 626 | A | 0.811 |
| 627 | E | 0.811 |
| 628 | G | 0.811 |
| 629 | D | 0.811 |
| 630 | V | 0.811 |
| 631 | G | 0.811 |

| | | |
|---|---|---|
| 632 | D | 0.811 |
| 633 | G | 0.811 |
| 634 | D | 0.811 |
| 635 | G | 0.811 |
| 636 | D | 0.811 |
| 637 | F | 0.811 |
| 638 | G | 0.811 |
| 639 | A | 0.811 |
| 640 | G | 0.811 |
| 641 | V | 2 |
| 642 | N | 2 |
| 643 | K | 1.5 |
| 644 | D | 2 |
| 645 | G | 1.5 |
| 646 | G | 1.5 |
| 647 | S | 0.811 |
| 648 | R | 1.5 |
| 649 | V | 0.811 |
| 650 | V | 1.5 |
| 651 | V | 1.5 |
| 652 | Q | 2 |
| 653 | M | 2 |
| 654 | E | 1.5 |
| 655 | E | 1.5 |
| 656 | V | 1.5 |
| 657 | A | 1.5 |
| 658 | R | 1.5 |
| 659 | T | 1.5 |
| 660 | V | 0 |
| 661 | N | 1.5 |
| 662 | V | 0.811 |
| 663 | V | 1.5 |
| 664 | M | 0.811 |
| 665 | V | 0.811 |
| 666 | L | 0.811 |
| 667 | V | 1.5 |
| 668 | P | 0 |
| 669 | L | 1.5 |
| 670 | L | 1 |
| 671 | L | 0 |
| 672 | L | 1.5 |
| 673 | L | 0 |
| 674 | C | 0.811 |

| | | |
|---|---|---|
| 675 | V | 1.5 |
| 676 | L | 0 |
| 677 | G | 1.5 |
| 678 | L | 0.811 |
| 679 | T | 1.5 |
| 680 | Y | 0.811 |
| 681 | A | 1 |
| 682 | L | 1.5 |
| 683 | V | 0.811 |
| 684 | Q | 0.811 |
| 685 | M | 0.811 |
| 686 | Q | 1.5 |
| 687 | R | 0.811 |
| 688 | K | 0.811 |
| 689 | G | 0.811 |
| 690 | A | 1.5 |
| 691 | P | 0 |
| 692 | R | 0.811 |
| 693 | V | 2 |
| 694 | L | 1.5 |
| 695 | L | 0.811 |
| 696 | Y | 0 |
| 697 | C | 0.811 |
| 698 | K | 0.811 |
| 699 | R | 0 |
| 700 | S | 0.811 |
| 701 | L | 1.5 |
| 702 | Q | 0.811 |
| 703 | E | 0.811 |
| 704 | W | 0.811 |
| 705 | V | 0 |

#### c. Equations

$$H = -\sum_{i=1}^{M} P_i \log_2 P_i$$

Equation 1. Shannon Entropy. For a multiple sequence alignment, $P_i$ is the fraction of residues for amino acid i, and M is the total number of amino acid types. H, Shannon Entropy, ranges from 0, where the same residue is present in that position for all sequences, to 4.332, where all 20 residues would be equally represented in that position. Residues with a Shannon Entropy greater than or equal to 2 are considered variable, while residues with values less than 2 are conserved, and residues with values less than or equal to 1 are highly conserved (Shannon)
